# Regulation of multiple dimeric states of E-cadherin by adhesion activating antibodies revealed through Cryo-EM and X-ray crystallography

**DOI:** 10.1101/2022.03.21.484361

**Authors:** Allison Maker, Madison Bolejack, Leslayann Schecterson, Brad Hammerson, Jan Abendroth, Tom E Edwards, Bart Staker, Peter J Myler, Barry M Gumbiner

## Abstract

E-cadherin adhesion is regulated at the cell surface, a process that can be replicated by activating antibodies. We use cryo-EM and X-ray crystallography to examine functional states of the cadherin adhesive dimer. This dimer is mediated by N-terminal beta strand-swapping involving Trp2, and forms via a different transient X-dimer intermediate. X-dimers are observed in cryo-EM along with monomers and strand-swap dimers, indicating that X-dimers form stable interactions. A novel EC4-mediated dimer was also observed. Activating Fab binding caused no gross structural changes in E- cadherin monomers but can facilitate strand swapping. Moreover, activating Fab binding is incompatible with the formation of the X-dimer. Both cryo-EM and X-ray crystallography reveal a distinctive twisted strand-swap dimer conformation caused by an outward shift in the N-terminal beta strand that may represent a strengthened state. Thus, regulation of adhesion involves changes in cadherin dimer configurations.

## Introduction

E-cadherin is a cell-cell adhesion protein^1–5^ that forms adherens junctions between epithelial cells. Dynamic regulation of cadherins at the cell surface makes them vital to proper tissue morphogenesis^2, 4, 6^ and is implicated in barrier function during inflammation^7, 8^ and in metastatic cancer^9–16^ when junctions are dysregulated.

A number of functional monoclonal antibodies (mAbs) to human E-cadherin extracellular domains have been identified that can activate, block adhesion, or distinguish between activation states of cell adhesion ^17, 18^. The group of activating mAbs has broad potential for therapeutic use: E-cadherin activating antibodies have been shown to decrease the number of metastatic nodules in mouse models of breast cancer^19, 20^ and can also decrease the disruption of barrier function and inflammation in mouse models of inflammatory bowel disease^8^. Other than the general location of epitopes on the E-cadherin extracellular domain^17^, the structural mechanism of activation resulting from these antibodies is as yet unknown. Understanding how these antibodies modulate E-cadherin function at the structural level would provide important insights into the mechanisms regulating the adhesive bond and have implications for development of potential therapeutics.

Although the mechanism underlying regulation of the cadherin adhesive bond at the cell surface is not well understood, a body of structural knowledge exists about the pathway underlying adhesive bond formation by individual cadherins. E-cadherin is a Type I classical cadherin, with 5 extracellular cadherin (EC) repeat domains with calcium binding sites between each, followed by a linker region, a single-pass alpha helical transmembrane domain, and a cytoplasmic tail^3, 21, 22^ complexed with ⍺-, β-, and p120-catenin linking the cadherin to the cytoskeleton^23, 24^. Cell adhesion is thought to occur through *trans* dimers between cadherins on opposing cells. *Cis* interactions between cadherins on the same cell have also been proposed to occur, forming a lattice proposed to form the adherens junction^22, 25, 26^, but mutations that block the cis interaction do not interfere with either cell adhesion^22^ or adherens junction formation^27^, and catenins^28^ and the transmembrane domain may also be involved^29^ in adherens junction assembly; thus the functional role of the cis dimer is unclear.

The stable final form of the *trans* interaction is thought to be mediated by the “strand-swap” dimer, named because it is mediated by the N-terminal beta strand in EC1 participating in a domain swap with the similar strand in the opposing cadherin EC1^22, 30–33^. The Trp2 residue of that strand in the monomer leaves a hydrophobic pocket in its own EC1 to enter the hydrophobic pocket of the opposing EC1 to form the dimer. The initial encounter complex during adhesive bond formation is thought to be via an intermediate in the monomer to strand-swap dimer transition, known as the X-dimer^31, 33–35^. This dimer is formed from an interface between EC1s, including a vital salt bridge between K14 and D138^31^ in the opposing EC1. This transition state brings the beta strands and Trp2 residues in proximity to each other, creating a favorable environment for the strand-swapping to take place. The Trp2 residues in the X-dimer are thought to flip to form a strand-swapped X-dimer^33^, and then this extends into the full strand-swap dimer^31–33^. Blocking the necessary salt bridge by mutating either K14 of D138 blocks the X intermediate, and these mutants are completely defective in cell adhesion.

It is important to note that all structures of the X-dimer to date have been of strand-swap deficient mutants; as of yet, this complex has not been observed in wild-type (WT) cadherins. As such, the X-dimer is thought to be a low-affinity state, as exhibited by the mutants^31, 33^, but it is difficult to know the prevalence of wild-type X- or strand-swap-X-dimers without observing them in solution.

In this work, we use cryo-EM and X-ray crystallography to explore the nature of E-cadherin dimer formation, as well as examine how dimerization is impacted by the binding of functional antibodies. Cryo-EM provides a way to observe dimer forms in solution in equilibrium without constraints imposed by crystallization or crystal packing. The activating and other antibodies bound to E-cadherin offer a means to examine how dimer forms are influenced by functional perturbation of the adhesive state, providing insights into possible mechanisms for enhancing E-cadherin adhesion.

## Results

### Multiple E-cadherin dimer conformations revealed by cryo-EM

In order to visualize cadherins in their most biologically relevant state, full-length human E-cadherin was purified and inserted into nanodiscs^36, 37^, preserving both the transmembrane and cytoplasmic domains along with the extracellular domain. Samples were then vitrified in 1 mM Ca^2+^-containing buffer and examined with cryo-EM. Some samples were also prepared with fully reconstituted E-cadherin-catenin protein complexes in nanodiscs, as described in a recent methodical study^38^. In the cadherin-catenin datasets, only E-cadherin and Fab were resolved and the catenins were never visible in the micrographs, indicating that they may have dissociated during freezing. No observable structural differences were noted between samples prepared with cadherin- only nanodiscs and cadherin-catenin nanodiscs (not shown), but because the catenins were likely dissociated, no conclusions about their effects, or lack thereof, can be drawn.

Calculation of 2D class averages of wild-type E-cadherin revealed that the nanodiscs were averaged out, presumably due to a flexible region between the extracellular and transmembrane domains. We confirmed the presence of nanodiscs with size exclusion chromatography (SEC) and negative stain EM (Extended Data Figure 1); nanodiscs alone were also visible in cryo-EM 2D averages (not shown).

Extracellular domains were noticeably rigid, producing distinct class averages (Figure 1C-E). We also note that there appeared to be only one cadherin per disc; we did not observe any *cis* interactions between cadherins that appeared to emerge from the same patch of lipids.

**Figure 1.**
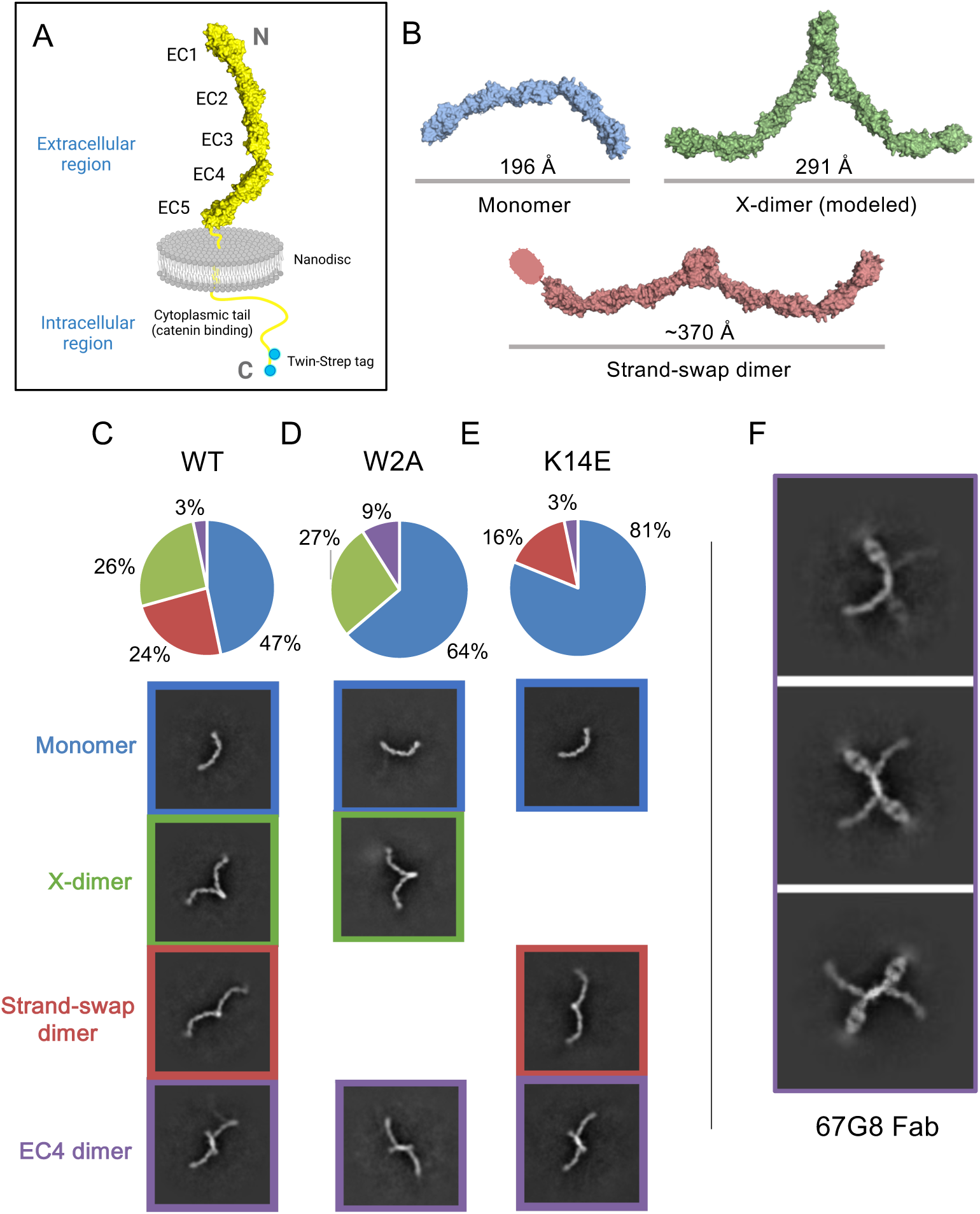
Cryo-EM 2D class averages of E-cadherin reveal monomers, X-dimers, and strand-swap dimer, as well as other novel dimer conformations. (A) Schematic of full protein used in this study. (B) Known and theoretical dimer conformations of E-cadherin. Monomer and strand-swap dimer: PDB 3Q2V. X-dimer created with alignment: PDB 3LNH, 3Q2V. (C) Class averages of WT full-length hE-cadherin include monomers, X- dimers, strand-swap dimers, and novel EC4 dimers. (D) Class averages of W2A full- length hE-cadherin include monomers, X-dimers, and EC4 dimers. (E) Class averages of K14E full-length hE-cadherin include monomers, strand-swap dimers, and EC4 dimers. (F) 67G8 EC5-binding Fab bound to FL-hE-cadherin indicates that novel dimers are indeed EC4-mediated.

2D class averages revealed a range of dimer conformations formed from the rigid cadherin monomers, including, as expected, structures similar to the strand-swap dimers observed in X-ray crystals, but also what appear to be X-dimers (Figure 1C). As X-dimers have previously only been observed in crystal structures of mutated EC1-2 fragments, we created a model of the full-length X-dimer by aligning the EC1-2 X-dimer crystal structure (PDB 4ZT1) to mouse E-cadherin EC1-5 monomers obtained from the crystal structure (PDB 3Q2V). This creates a structure much like we see in our 2D averages, a compacted dimer with a diameter of ∼290 !, compared to ∼370 ! for the more extended strand-swap dimer (Figure 1B). Although proportions varied somewhat between grid conditions, we saw what appeared to be a similar number of particles in both strand-swap and X-dimer conformations (Figure 1C) in repeated datasets. We repeated each mutant in the same conditions 2 times; Figure 1 indicates results from one dataset.

To verify that the extended and more compact structures are indeed strand-swap and X-dimers, respectively, we introduced mutations into E-cadherin that are known to interfere with dimer formation. Strand-swap dimers are disrupted by the W2A mutation, which eliminates the strand-swap binding^31^. X-dimers are blocked through the K14E mutation, which inverts the charge of a salt bridge in the dimer interface^31^. We observed that W2A E-cadherin 2D averages only exhibit monomer and compact X-dimer conformations (Figure 1D), whereas K14E E-cadherin only forms extended dimers and monomers (Figure 1E). Thus, these mutants support our hypothesis that the compacted dimer is an X-dimer and the extended dimer is a strand-swap dimer.

The presence of the X-dimer in these samples was unexpected because the X-dimer is thought to be a low affinity short-lived transition state. One possibility is that we may be observing combined strand-swapped X-dimers, which have been postulated to occur using molecular dynamics simulations of E-cadherin^39^ and have been observed in P- cadherin mutants^33^. This conformation may be more stable than unswapped X-dimers. However, when we introduce mutations in the Trp2 residue preventing strand-swapping, a high proportion of compact X-dimers are still visible. Therefore, the observed X-dimers must include a substantial fraction of non-strand-swapped X-dimers.

In addition to the two *trans* dimers known from crystallography data, we also observed a novel dimer that appears to be mediated by an interaction between the EC4 domains of two opposing cadherins. This dimer was seen in a much smaller but significant percentage of particles over a variety of grid and sample conditions. To confirm the location of the interaction site, we used EC5 binding Fab 67G8 as a marker to determine the E-cadherin orientation in the 2D averages (Figure 1F). The Fab’s location close to the dimer interface indicates that this is indeed an EC4-EC4 association. This suggests the possibility that EC4 dimerization may have a role in cadherin function, but it is difficult to discern its impact from this structural information alone.

### Effects of functional antibody binding on monomeric E-cadherin

Previous work demonstrated the dramatic effects of functional antibodies to hE-cadherin on cell adhesion, particularly activating antibodies on cells^17–19^, as well as in animal models^19, 20^. However, little is understood about the biochemical or structural mechanism of these activating antibodies. Based on our cryo-EM observations of E-cadherin structures, we sought to determine how the binding of activating antibodies affects the conformational landscape of E-cadherin monomers and dimers.

By mixing E-cadherin nanodiscs with Fabs, we were able to reliably determine cryo-EM 3D reconstructions for several functional Fabs bound to E-cadherin. We compare structures of two activating antibodies (59D2 and 19A11), a control neutral antibody that has no effect on adhesion (46H7), and an adhesion blocking antibody (67G8) (Figure 2)^17^. The resolution of each of these ranged from 4.7-6.2 !, providing unambiguous docking for each Fab with the cadherin. As noted in previous epitope mapping work^17^, the two activating antibodies bind near the same site, on the opposite side of EC1 from the adhesive Trp2 strand (Figure 2A,B,D). The control neutral antibody 46H7 binds the outer curve of EC3 (Figure 2A, C). Blocking antibody 67G8 binds the end of EC5 (Figure 2A, E).

**Figure 2.**
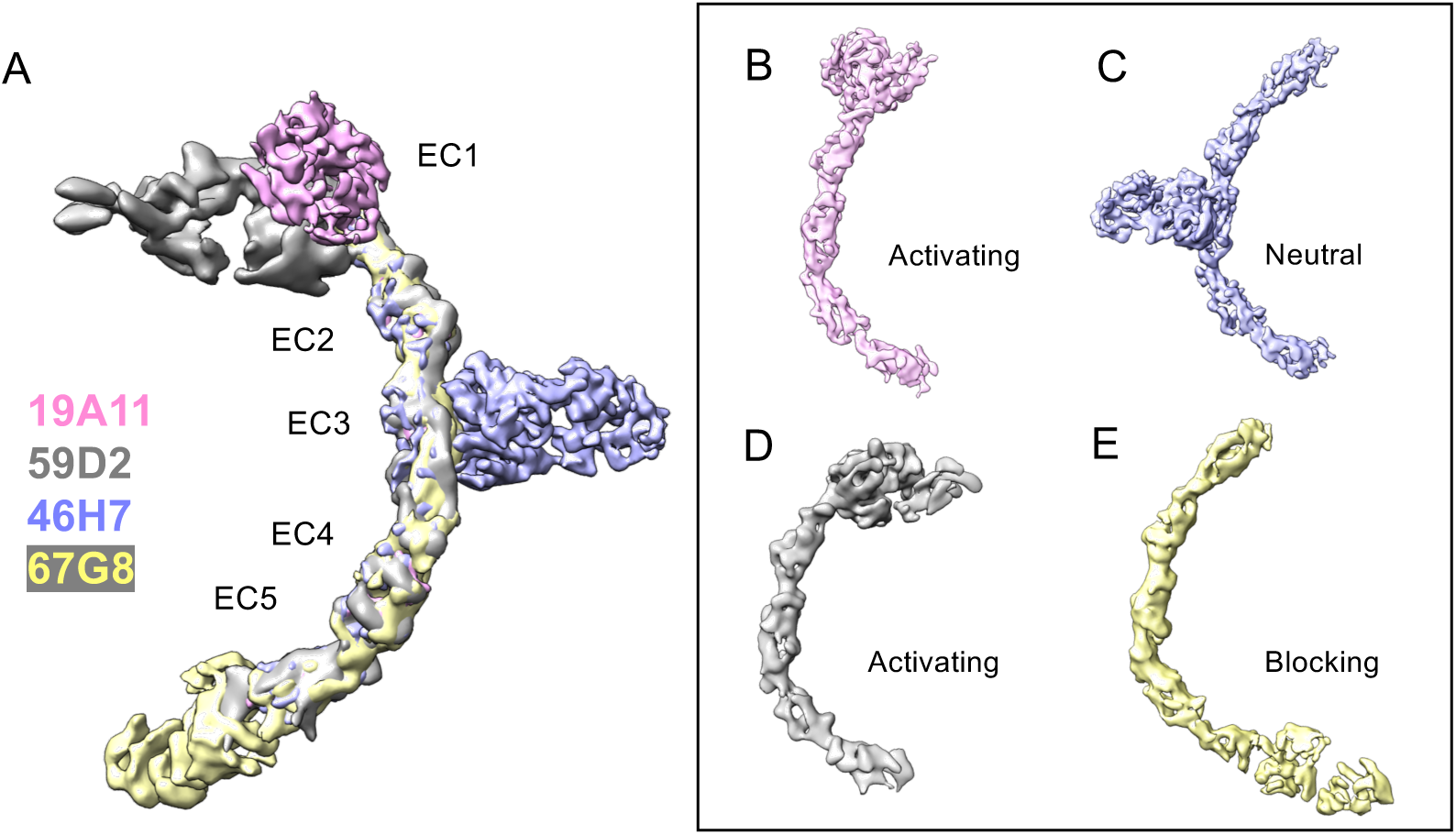
Cryo-Electron microscopy reconstructions show that monomeric structure of E-cadherin is not dramatically affected by activating Fab binding. (A) Overlay of all structures (B) hEC1-5 with activating Fab 19A11 (C) hEC1-5 with neutral Fab 46H7 (D) hEC1-5 with activating Fab 59D2 (E) hEC1-5 with inhibitory Fab 67G8.

Although the activating antibodies have been reported to have allosteric effects on adhesion, none of them induce any notable large-scale conformational changes in E-cadherin monomers (Figure 2A); nor did combinations of multiple antibodies (not shown). Compared to the crystal structure of dimeric mouse E-cadherin (PDB 3Q2V) there is a subtle curvature difference, particularly in the more kinked Ca^2+^ binding site between EC3-4 (not shown), but this curvature difference was observed for all the antibodies, including the neutral control. This small difference is unlikely due to Fab binding and may instead be due to reduced forces on the protein outside of crystal packing and in its monomeric form. We also note that cryoSPARC 3D variability analysis of all structures suggests some potential flexibility between EC3-4.

The 5-6 ! resolution of all these structures limits our ability to detect more atomic level effects of antibody binding to monomeric E-cadherin that might be important for the mechanisms of their effects on adhesion. Nonetheless, in the structures with bound activating Fabs, there is a notable lack of density in the middle of EC1, and a corresponding increase in density extending out from where the N-terminal Trp2 strand would extend (Extended Data Figure 2). This change in densities is not evident in the structure with neutral antibody 46H7 bound. This change in density with activating Fab may be significant because the model in the literature for the monomeric state of E-cadherin has the Trp2 forming an intramolecular bond, filling the hydrophobic pocket in its own subunit, and the extrusion of the Trp2 leads to the intermolecular strand-swap dimer that underlies adhesive bond formation. Although not high resolution, our observation raises the possibility that activating Fabs act to destabilize the monomeric state of E-cadherin. Increasing conformational strain in the N-terminal strand in the monomer through the E11D mutation was shown to increase dimer affinity in previous studies^40^, and activating Fabs could be doing something similar.

### Activating Fabs are compatible with strand-swap dimers but not X-dimers

When activating Fabs are bound to E-cadherin, a significant number of strand-swap dimer particles are observed in the 2D averages in the cryo-EM datasets, but we never observed X-dimers. In the case of 19A11 bound to WT E-cadherin both strand-swap dimers and monomeric E-cadherin were present, but not X-dimers (Figure 3A). The dimers are strand-swapped dimers because 19A11 bound to the E-cadherin W2A mutant protein revealed only monomeric cadherin (Figure 3B). We also did not observe X-dimers in any E-cadherin/59D2 Fab datasets (not shown). Overall, it is very clear that activating Fabs can form complexes with strand-swapped dimers, but not with X-dimers.

**Figure 3.**
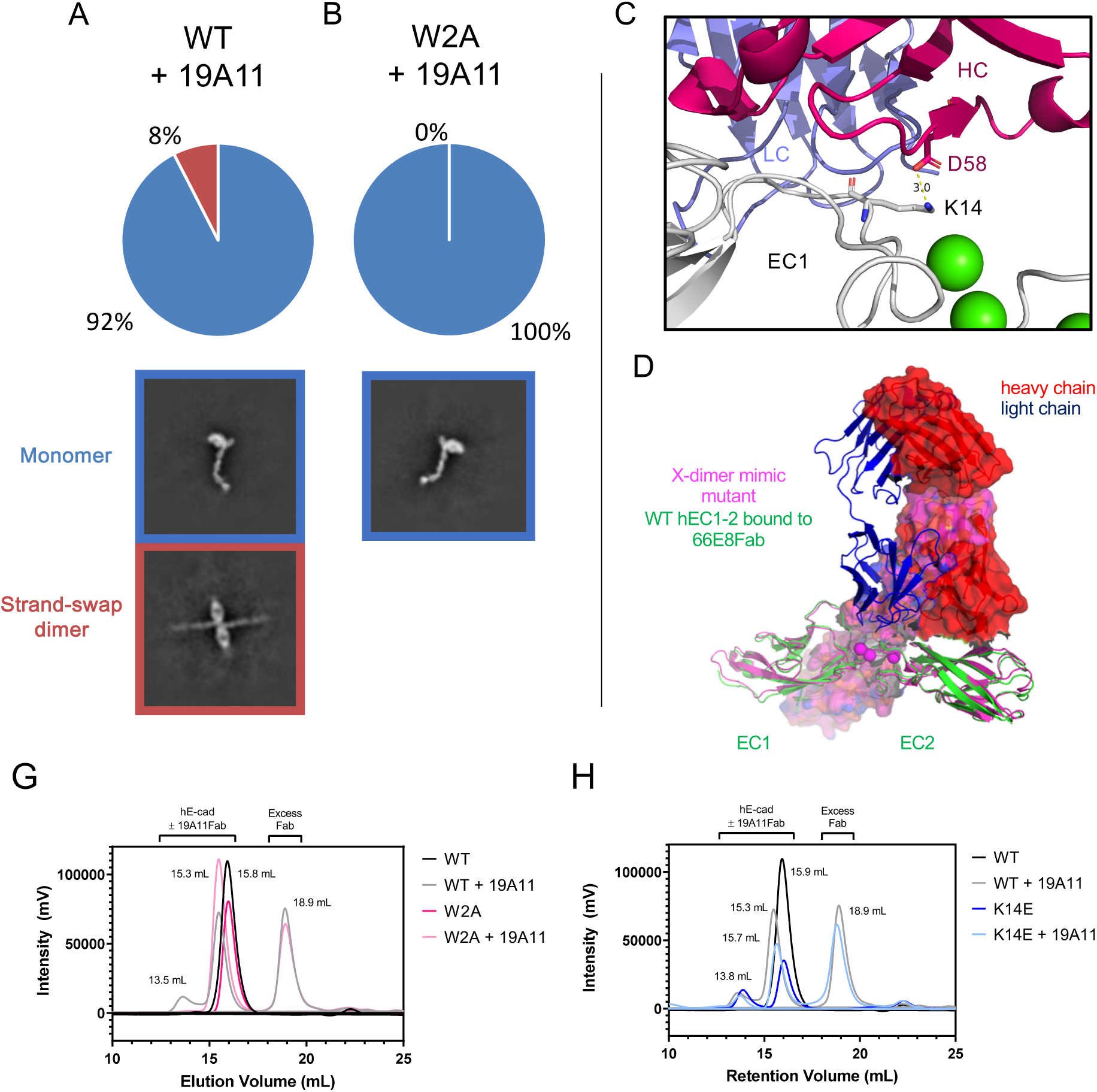
19A11 activating antibody bound to E-cadherin is not seen to co-exist with X- dimer intermediate. (A) Class averages of 19A11 bound to WT full-length hE-cadherin include monomers and strand-swap dimers. (B) Class averages of 19A11 bound to W2A full-length hE-cadherin include only monomers. (C) K14 of hE-cadherin forms a salt bridge with D58 on the 19A11 heavy chain. (PDB 6CXY) (D) The heavy chain of the 66E8 activating Fab would have a massive steric clash with the theoretical X-dimer, indicated in magenta (PDB 4ZT1). (G) In SEC, 19A11Fab binding to hEC1-5 triggers the formation of a strand-swap dimer peak, blocked by the W2A mutation. (H) 19A11 Fab bound to hEC1-5 WT shows an analogous peak pattern to hEC1-5 K14E, the X- dimer blocking mutant.

X-ray crystallographic data also demonstrate an incompatibility between the X-dimer state and activating Fab binding to E-cadherin. We obtained X-ray structures of two activating Fabs bound E-cadherin. Activating Fab 66E8 was crystallized with the hEC1-2 fragment (Figure 3D, Extended Data Figure 4), and 19A11 Fab was crystallized with either hEC1-2 (Figure 3C, Extended Data Figure 3) or the full hEC1-5 ectodomain (Figure 4). All Fab-bound E-cadherins crystallized into strand-swapped dimer structures. These crystallographic data also demonstrate an incompatibility between the X-dimer state and activating antibody binding to E-cadherin. Both crystal structures of 19A11 bound to E-cadherin (EC1-2 and EC1-5) show that Fab binding involves a salt bridge between the heavy chain of 19A11 and K14 on E-cadherin (Figure 3C). As the K14- D138 E-cadherin dimer salt bridge is necessary for X-dimer formation, and the affinity of 19A11 to E-cadherin at ∼6.5 nM (Extended Data Figure 5A,F) is on the order of 10^4^ times stronger than the affinity of any dimer of WT E-cadherin (∼100 uM^31^), it is unlikely that the X-dimer would supersede 19A11 binding. The crystal structure of 66E8 Fab, another activating antibody, bound to hEC1-2 (Extended Data Figure 4), suggests that the bound Fab would cause a complete steric clash with X-dimer formation (Figure 3D). Although the affinity of 66E8 is weaker than 19A11 at ∼100 nM (Extended Data Figure 5E,F), it still surpasses that of cadherin dimers. Thus, it appears that two different activating antibodies structurally interfere with the X-dimer.

**Figure 4.**
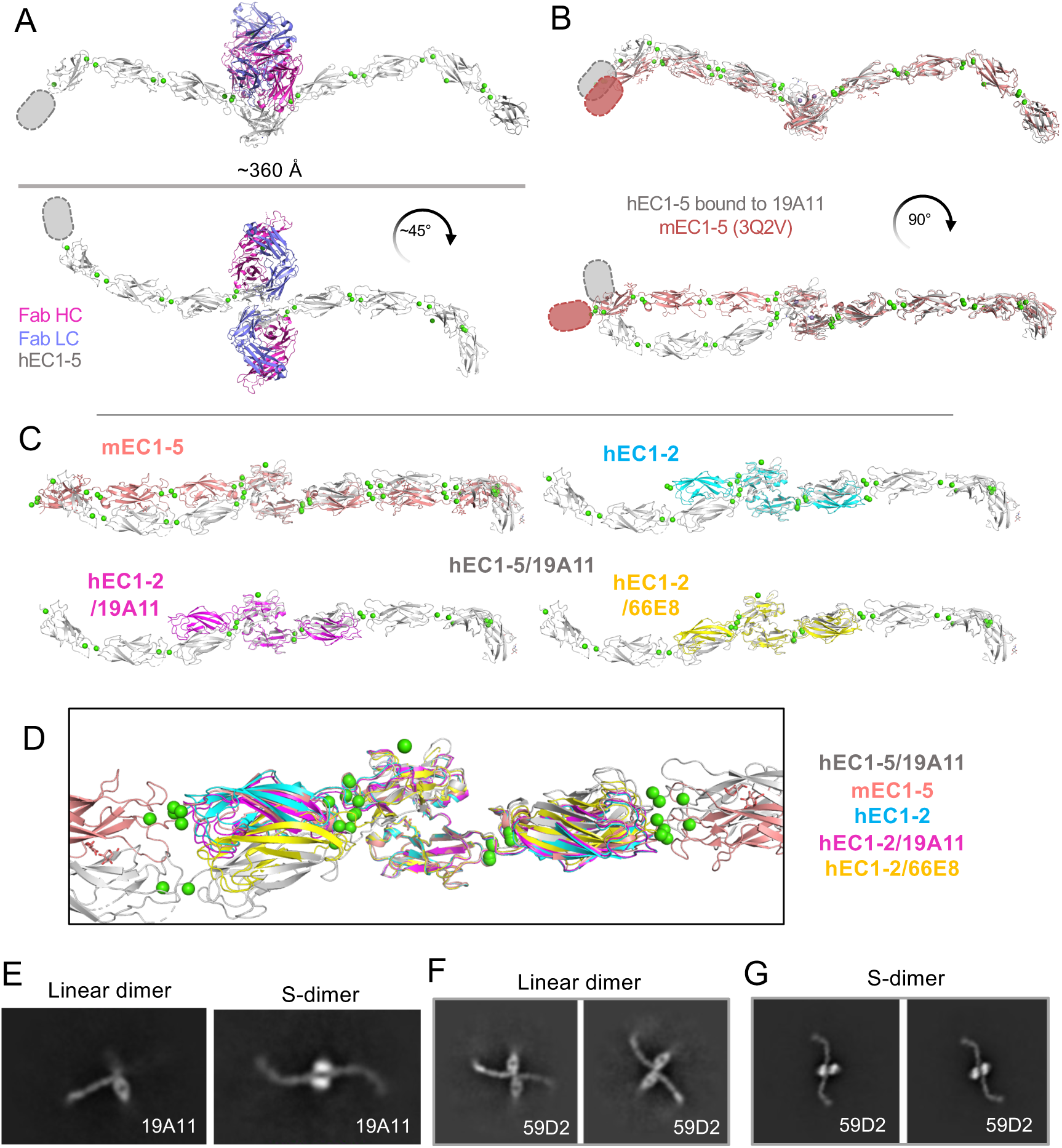
Activating antibody reveals a novel, tightened “S” dimer conformation in human E-cadherin, influenced by Trp2 positioning as well as a EC1-2 Ca^2+^ site bend. (A) Overall crystal structure of hEC1-5/19A11Fab dimers, highlighting twisted conformation. Missing EC5 density indicated with ovals. (B) hEC1-5 bound to 19A11 with one monomer aligned with mouse EC1-5 (PDB 3q2v); Fabs removed for clarity. (C) Comparison of hEC1-5/19A11 dimer orientation with other E-cadherin structures. EC1 of the right monomer was aligned on each. (D) All EC1 alignment dimer structures overlaid. (E) Both straight and twisted strand-swap dimers seen in dataset of 19A11Fab bound to the complete cadherin-catenin complex. (F) Class averages of activating Fab 59D2 with hE-cadherin indicate canonical strand-swap dimer (G) Class averages of 59D2 with full cadherin-catenin complex show the twisted strand-swap conformation.

All these data showing incompatibility of activating Fab binding with X-dimer formation is difficult to reconcile with the proposed role of the X-dimer as a required transition state intermediate towards formation of strand-swap adhesive dimers.

Mutations that interfere with the formation of the X-dimer prevent cadherin adhesion in cell models^19, 31^. One possibility is that activating antibodies could potentially allow skipping of the intermediate state. However, Petrova et al.^19^ found that 19A11 activating antibodies did not rescue adhesion by K14E - E-cadherin mutants in cell adhesion assays. We repeated the experiment with multiple activating antibodies, including 66E8 and 59D2 (Extended Data Figure 8A), and found that none of them were able to rescue the X-dimer blocking mutation in cell adhesion. Thus, either the X-dimer intermediate is still required, or the K14 residue has other roles in adhesion.

In cryo-EM, we did not observe an increase in the fraction of strand-swapped dimers in the presence of activating Fabs. For all antibodies, including the control neutral Fab, a small decrease in the proportion of dimers was observed, and we suspect that changes in protein concentrations in ice rather than any effects of the Fabs may have been responsible.

Although we did not observe an increase in strand-swapped dimers in nanodisc cadherin preparations by cryo-EM, we also examined whether activating antibody 19A11 exhibited any biochemical effects on the formation of strand-swap dimers of soluble non-membrane associated cadherins. When Fabs were incubated in excess with soluble full E-cadherin ectodomains (hEC1-5) and analyzed by size exclusion chromatography (SEC) 19A11 Fabs increased E-cadherin dimerization (Figure 3G). All other antibodies appeared to form complexes only with monomeric hEC1-5 at readily workable concentrations (Extended Data Figure 6). Dimers are represented by an early peak in the SEC trace at ∼13.5 mL. 19A11 Fab bound to the hEC1-5 W2A strand-swap incapable mutant, but the early peak was no longer evident (Figure 3G), demonstrating that the early peak in the WT trace was a strand-swap dimer. The X-dimer blocking K14E – E-cadherin mutant protein alone also eluted with a separated strand-swap dimer and monomer peak pattern (Figure 3H), consistent with previous studies showing that it can still form strand-swap dimers at equilibrium^31^. Although K14 is part of the epitope, 19A11 was also shown by SEC to be able to bind the K14E mutant (Extended Data Figure 6F), although likely more weakly, but it did not affect monomer/dimer proportions (Figure 3H). These data suggest that 19A11 induces the formation of strand-swap dimers in solution, similar to the effects of the K14E X-dimer blocking mutation.

### Activating antibodies induce changes in the structure of the strand-swap dimer

Examination of the strand-swapped dimer structures in crystal structures of activating Fab bound E-cadherin and comparisons with observations of Fab-bound dimers seen in cryo-EM revealed two very different strand-swap dimer conformations across the same dimer interface (Figure 4). Most notably, 19A11 Fab bound to EC1-5 (PDB 7STZ) (Figure 4A) crystallized in a different conformation than that of either E-cadherin alone (PDB 2O72) or of 19A11 Fab bound to EC1-2 (PDB 6CXY) (Figure 4B- D). From a quaternary structure standpoint, crystal structures of Type I classical cadherins form a W shape when observed from the side and appear linear when observed from the top. The structure of hEC1-2/19A11 (PDB 6CXY) formed an analogous conformation to the linear conformation of the mouse E-cadherin dimer (PDB 3Q2V) (Figure 4C,D). However, 19A11 in complex with the full E-cadherin ectodomain, hEC1-5/19A11, forms a twisted conformation when viewed from the top, resembling an “S” – henceforth referred to as the S-dimer (Figure 4A, B). The diameter of this S conformation is ∼360 Å, compared to the linear strand-swap diameter of 370 Å, revealing a slightly compacted structure.

Notably, in one cryo-EM dataset of 19A11 Fab bound to E-cadherin, we noticed 2D class averages for both dimeric conformations (linear and S), as shown by the dimer shape and degree of Fab protrusion (Figure 4E). Additionally, when examining another activating Fab, 59D2, we observed both conformations in two separate datasets of 59D2/hE-cadherin and 59D2/full-cadherin catenin complexes (Figure 4F,G). (66E8 activating Fab tended to self-associate, so we were unable to assess dimeric states of E-cadherin bound to this antibody with EM). Both conformations can be seen when the same activating Fab is bound. Importantly, the “S” conformation was only observed when activating Fabs were bound to E-cadherin, not with the neutral or blocking Fabs. The fact this conformation was seen in solution with two different activating antibodies in addition to the 19A11Fab-EC1-5 crystal structure lends credence to it being biologically relevant and not a crystal packing artifact.

Examination of the molecular details of the dimer interaction in the S conformation indicate that there is a difference in the angle between EC1 domains at the strand-swap interface compared to other hE-cadherin crystal structures (Figure 4C, D). There is also a bend between EC1 and EC2 at the calcium binding site that is most prominent when compared to mouse EC1-5. This increases the twist in the dimer in the overall structure in addition to the angle shift between EC1s. Interestingly, the degree of this bend appears to correlate with EC1 dimer angle, indicating the two changes may be linked.

The only significant conformational change in EC1 between the linear strand-swap and S-dimer is a symmetric inward shift of the first four N-terminal residues (DWVI) of both monomers, with the shift most notable in the Trp2 residue (Figure 5). In fact, although hEC1-5/19A11 shows by far the most pronounced inward shift in known E-cadherin strand-swap structures, the crystal structure hEC1-2 bound to another activating Fab, 66E8, also exhibits this inward N-terminal shift (Figure 5C), as well as the aforementioned bend at the Ca^2+^ site (Figure 4F). This Trp2 shift appears to be in the same plane, with no rotation (Figure 5A,D). Interestingly, there appear to be no modifications of the hydrophobic pocket in which the Trp2 binds with this linear shift; the opposing protomer Trp2 fits into an identical position in the first protomer pocket regardless of Trp2 shift (Figure 5D).

**Figure 5.**
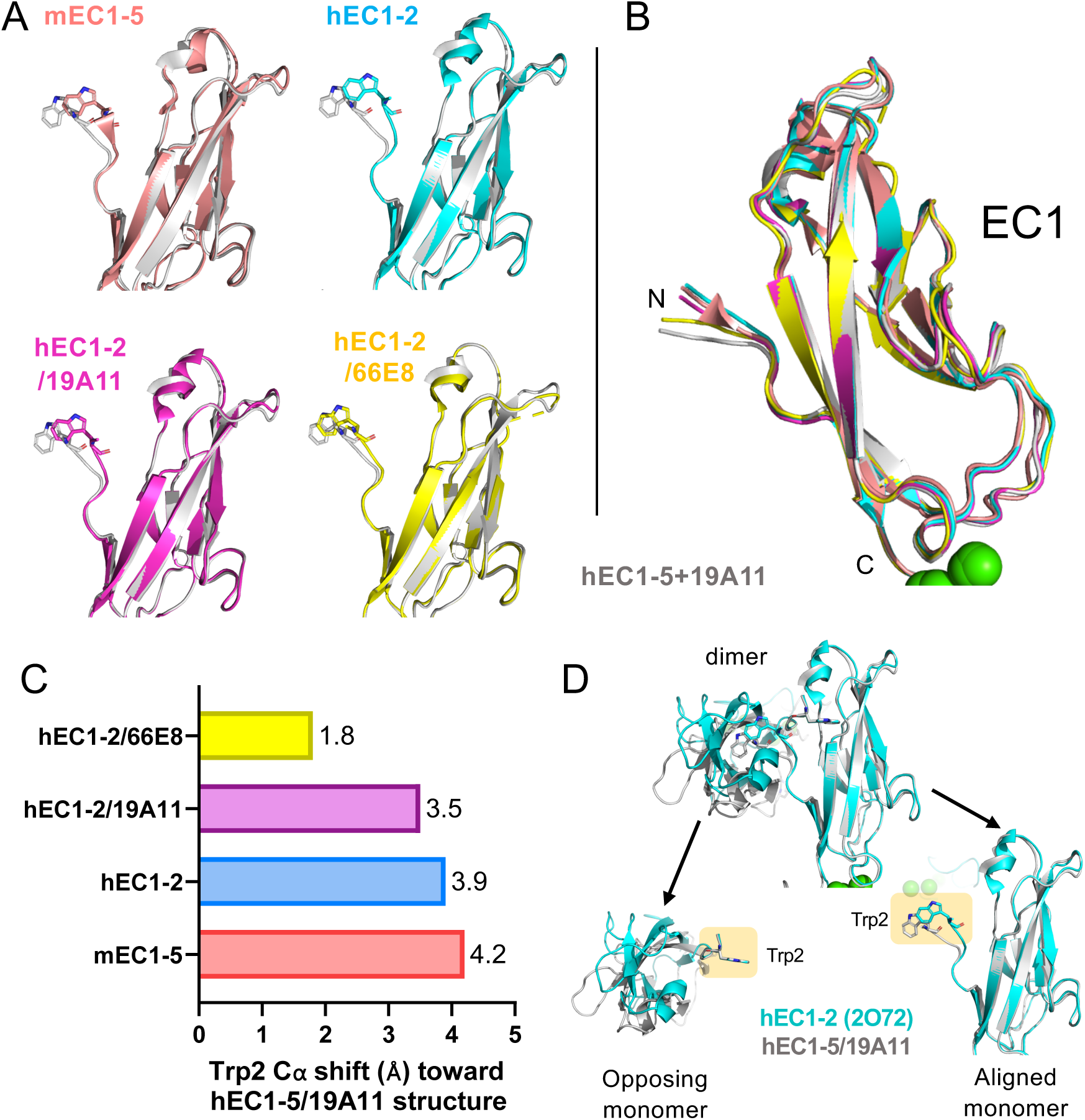
Comparison of Trp2 position with other E-cadherin structures. (A) Individual EC1 structures compared to 19A11/hEC1-5. The Trp2 residue is highlighted. Opposing dimers, as well as Fabs in Fab-bound structures removed for clarity. (B) All EC1 structures overlaid. (C) Inward shift of Trp2 C⍺ compared to hEC1-5/19A11 structure. (D) Comparison of hEC1-5/19A11 strand-swap and hEC1-2 strand swap. Monomer 2 is aligned in each. The Trp in monomer 1 does not move, but the EC1 shifts, and vice- versa in monomer 2.

## Discussion

This work describes the regulation of the E-cadherin adhesive bond as a multistate process, involving a variety of conformations, and provides potential mechanisms for how the bond is regulated by activating antibodies. Previous research has provided evidence for two cadherin trans-dimer states using X-ray crystallography and by altering the cadherin with mutations^31, 33, 35, 40^ or blocking antibodies^41^. Using cryo-EM to examine cadherins in solution, we are able to discern both of these dimer conformations in wild-type E-cadherins, as well previously unreported dimer conformations, including an EC4-mediated dimer, and an “S” shaped adhesive strand-swap dimer that was observed only when bound to activating antibodies.

This is the first visual evidence of X-dimers forming with WT cadherins, with a mixture of X- and strand-swap dimers occurring together, which is surprising because the X-dimer is thought to represent a low affinity and very transient intermediate. This indicates that X-dimers are much more stable than previously recognized at relatively low concentrations in solution. All previous measurements indicating a very low affinity of X-dimer formation have been done using strand-swap incompetent W2A mutants.

When the affinity of WT E-cadherin dimers is measured, there is no way of knowing which dimer conformation is being analyzed without structural information. It is possible that some amount of the X-dimers we observe in cryo-EM represent a “strand-swapped X-dimer” that has been proposed to be a secondary intermediate between X- and strand-swap dimers^33^, in which the cadherins are in the X-dimer conformation, and the Trp2s are strand-swapped simultaneously. However, the large number of X-dimers that we observed with E-cadherin harboring the W2A mutation cannot be strand-swapped, and therefore must represent a larger proportion than expected of unswapped X-dimers. Although we cannot determine actual affinities from our cryo-EM experiments, these findings show that the X-dimer exists as a somewhat stable conformation in solution which does not necessarily advance to forming the strand-swap dimer conformation.

This raises the possibility that both strand-swapped and X-dimers can exist in equilibrium on the cell surface at the adhesive interface.

It is challenging to reconcile our findings from both cryo-EM and Xray crystallography that activating Fab binding to E-cadherin is structurally incompatible with X-dimer formation with earlier findings that the X-dimer is a requisite intermediate. It may be necessary to revisit this model for the pathway of adhesive bond formation.

However, one possibility is that activating Fabs act by preventing the reverse reaction from strand-swap to X-dimer after the strand-swap adhesive dimer has already formed, preventing adhesive bond dissociation. This is the most likely explanation for the 66E8 activating antibody which has a large steric clash with X-dimer, and would be consistent with the model of adherens junction dissociation via the X-dimer proposed by Hong et al^42^. Another intriguing possibility suggested by the nature of 19A11 Fab-cadherin structure is that Fab binding enhances the transition from X-dimer to strand-swap dimer. 19A11 Fab forms its own salt bridge with K14, a residue vital to formation of the putative X-dimer intermediate. At first, this idea appears to be counterintuitive because mutations blocking the X-dimer block E-cadherin adhesion in cells. However, 19A11 Fab is still capable of binding the K14E mutant of E-cadherin as observed in SEC (Figure 3H, S7F), indicating that this residue is not necessary for binding even though it is part of the full epitope. Therefore, 19A11 could bind initially when the K14 residue is participating in the X-dimer dimer interface and then “steal” the salt bridge, forcing it from the X-dimer into the strand-swap conformation. Our observation of X-dimers co- existing with strand-swap dimer in solution raises the possibility that a dynamic equilibrium with both states exists at cell-cell adhesion sites. Then activating antibodies could drive the dimer into the strand-swapped conformation, resulting in a strengthened state of adhesion since the strand-swap may be the more favored state under tension^43^.

The new twisted “S” dimer conformation that we observe in both cryo-EM and X- ray crystallography may also represent a further strengthened strand-swapped state, since it was observed only in the presence of activating antibodies. This conformation arises mainly from a shift in the first 4 N-terminal residues (DWVI) of the beta strand important in the monomer to dimer conversion. Vendome et al.^40^ emphasized the importance of beta strand instability leading toward E-cadherin strand-swap dimerization. The rigidity imposed by the calcium binding sites, primarily mediated by Glu11, which exists at the hinge point of the beta strand, contribute to “conformational strain” of the beta strand, promoting its expulsion during strand-swap binding. All 3 activating antibodies studied bind at or near the anchor points on the opposite side of the cadherin from the beta strand. 19A11 Fab binds the back side of EC1 including residues close to the Glu11 hinge point; 66E8 and 59D2 bind in the calcium binding region, all consistent with this model.

Finally, in addition to X-, linear strand-swap, and S-strand-swap dimers, other E-cadherin dimers may exist that are important for regulation of adhesion. We observe of a reproducible EC4-mediated dimer in cryo-EM (Figure 1C-F), although it is unknown whether this dimer is biologically relevant. The blocking antibody 67G8 bound to E-cadherin showed a high proportion of EC4 dimers (Figure 1F) suggesting that it could have some association with inhibiting adhesion. Furthermore, there are many indications in the literature that this region is important for controlling cell adhesion.

Several activation distinguishing antibodies bind at the EC3-4 boundary^17^. Also, many gastric cancer mutations are in the EC4-5 region of E-cadherin^11, 19^, and aberrant N-linked glycosylation of Asn554 (Asn 400 – mature protein) in EC4 has been linked to poorer gastric cancer outcomes and weakened adhesion^44^. Similarly, the *half-baked* mutation in the EC4 domain of E-cadherin disrupts morphogenesis of early zebrafish embryos^45^. Moreover, biophysical studies have found cadherins to undergo a distance dependent three-step unbinding process under force involving the EC3-4 domains as well as EC1-2^46^. This evidence for a functional role for EC4 suggests that the EC4 dimer could have a role in cadherin regulation and should be explored in future studies.

This study highlights the complexity of the landscape of E-cadherin *trans*-dimer states and the roles they play in adhesion regulation by activating antibodies. The effect of activating Fabs on the X-dimer raises the possibility that the canonical pathway from monomer to X-dimer to strand-swap dimer needs modification. Alternatively, the antibodies could act by binding to and dissociate existing X-dimers to induce adhesion or by preventing adhesive bond dissociation by preventing reversion to the X-dimer. In addition, more subtle and complex structural changes in the conformation of the strand-swap adhesive bond associated with activating Fab binding may modulate the intricate dynamic regulation of E-cadherin adhesive binding states.

## Materials and Methods

### Protein Expression and Purification

#### Full-length E-cadherin

Expression and purification were done following the protocol from our previous work reconstituting the cadherin-catenin complex^38^. We used the full sequence of human E-cadherin with the signal sequence and pro-domain deleted (Δ1- 154), an alternative CD33 signal sequence inserted (GMPLLLLLPLLWAGALA) before the N-terminal residue, and a Twin-Strep tag added after the C-terminal residue (SAWSHPQFEKGGGSGGGSGGGAWSHPQFEK*). This was cloned into pcDNA3.4 and transfected into Expi293 cells (ThermoFisher) with the ExpiFectamine 293 Expression Kit (ThermoFisher) according to standard protocols. Four days post-transfection, cells were spun down, and pellets were stored at -80°C until purification. The base buffer used for all purification steps is Strep Binding Buffer (BB): 50 mM Tris, 150 mM NaCl, pH 8.0. Upon purification, cell pellets were thawed on ice, then resuspended with 2x pellet volume of lysis buffer: BB + 1mM CaCl_2_ + 1% IGEPAL® CA- 630 (Sigma 56741) + 10 μL HALT protease inhibitor cocktail (ThermoFisher 78425)/mL total volume + 18.1 mL BioLock (IBA 2-0205-050)/mL pellet volume. Resuspended pellets were lysed gently rocking at 4°C for 45 minutes, then insoluble material was removed by spinning 25000xg for 15 min. Cleared supernatant was loaded into a StrepTactin XT gravity column (IBA) equilibrated in BB + 1mM CaCl_2_ + 1% IGEPAL® CA-630, then washed with BB + 1mM CaCl_2_ + 1% IGEPAL® CA-630, then BB + 0.02% lauryl maltose neopentyl glycol (LMNG) (Anatrace). Protein was eluted in BB + 1mM CaCl_2_ + 0.02% LMNG + 50 mM D-Biotin (IBA), then buffer exchanged into BB + 1mM CaCl_2_ + 0.02% LMNG with a PD-10 column and either flash frozen and stored at -80°C or immediately used. Protein quality was then assessed by SDS-PAGE and SEC using a Superose 6 10/300 GL (GE) column.

#### E-cadherin extracellular domains

We used residues 155-698 to encompass EC1-5 of the human E-cadherin extracellular domain. Similarly to full-length E-cadherin, the signal sequence and pro-domain were deleted (Δ1-154), an alternative CD33 signal sequence was added, and a Twin-Strep tag added after the C-terminal residue. E- cadherin used for BLI had an additional 8His tag (HHHHHHHH) after the TwinStrep tag. These constructs were cloned into pcDNA3.4 and transfected into Expi293 cells (ThermoFisher) with the ExpiFectamine 293 Expression Kit (ThermoFisher) according to standard protocols. If protein was to be used for crystallization, 5 "M kifunensine was added at time of transfection to limit glycosylation processing. As this protein was secreted into the medium, cells were spun down 4 days post-transfection, and supernatant was retained and 0.2 μM filtered. If protein was for crystallization, 500000 U Endo Hf (NEB) was added to the filtered supernatant and incubated for 1-2 days before purification to remove branched glycans. Cell culture supernatant was treated with 18.1μL/mL BioLock (IBA), 10x BB to 1x, and CaCl_2_ to 1 mM for 15 min to block biotin from binding the StrepTactin column and create favorable buffer conditions for column binding. Supernatant was then loaded into a StrepTactin XT gravity column (IBA) equilibrated in BB + 3mM CaCl_2_, then washed with BB+ 3mM CaCl_2_. Protein was eluted in BB + 3mM CaCl_2_ + 50 mM D-Biotin (IBA), then buffer exchanged back into BB + 3mM CaCl_2_ with a PD-10 column and flash frozen and stored at -80°C. Protein quality was assessed by SDS-PAGE and SEC using a Superose 6 10/300 GL (GE) column.

#### hE-cadherin EC1-2

We used residues 155-371 to encompass EC1-2 of the human E-cadherin extracellular domains. Similarly, to full-length E-cadherin, the signal sequence and pro-domain were deleted (Δ1-154). EC1-2 was expressed as a fusion protein by attaching 6x His tagged SMT3 to the N-terminus (PMID: 18467498). The EC1-2 construct was cloned into pET21a plasmid system and transformed into BL21 (DE3) competent cells (Novagen). Cultures were grown in autoinduction media (PMID: 15915565) overnight and harvested via centrifugation. Thawed bacterial pellets were lysed by sonication in 200 ml buffer containing 25 mM HEPES pH 7.0, 500 mM NaCl, 5% Glycerol, 0.5% CHAPS, 10mM Imidazole, 10 mM MgCl2, and 3 mM CaCl2. After sonication, the crude lysate is clarified with 2µl (250 units/ul) of Benzonase and incubated while mixing at room temperature for 45 minutes. The lysate is then clarified by centrifugation at 10,000 rev min−1 for 1 h using a Sorvall centrifuge (Thermo Scientific) followed by filtration via 0.45µm syringe filters. The clarified supernatant was then passed over a Ni-NTA His- Trap FF 5 ml column (GE Healthcare) which was pre-equilibrated with loading buffer composed of 25 mM HEPES pH 7.0, 500 mM NaCl, 5% Glycerol, 20 mM Imidazole, and 3 mM CaCl2. The column is washed with 20 column volumes (CV) of loading buffer and was eluted with loading buffer plus 500 mM imidazole in a linear gradient over 10 CV. Peak fractions, as determined by UV at 280 nm, are pooled and concentrated to 10mL. Pooled fractions are dialyzed overnight against 4 liters buffer containing 500mM NaCl, 25mM HEPES, 5% Glycerol, 3mM CaCl2 (SEC Buffer) with His-tagged Ulp1 protease added to cleave the 6xHis-SMT3 fusion protein at a ratio of 1 mg Ulp1 for 1000 mg protein. Dialysate is passed over a Ni-NTA His-Trap FF 5 ml column to remove 6xHis-SMT3 fusion protein and Ulp1. Flowthrough from the nickel column is concentrated to 5 ml and passed over a Superdex 75 SEC column (GE) equilibrated with SEC Buffer. The peak fractions were collected and analyzed for the presence of the protein of interest using SDS–PAGE. The peak fractions were pooled and concentrated using Macrosep 20 mL 10K MWCO protein concentrators (Pall). Aliquots were flash-frozen in liquid nitrogen and stored at −80°C until use for crystallization or in preparation of hE-cadherin EC1-2-Activating Fab complexes.

#### Fabs

Sequences coding for the heavy chain of Fab fragments were cloned into pcDNA3.4 with either a C-terminal 6His-tag or Twin-Strep tag sequences described above.

ExpiCHO cells (ThermoFisher) were transfected with the appropriate light chain and heavy chain encoding plasmids for each Fab following the ExpiFectamine CHO Transfection Kit (ThermoFisher) high titer protocol. Purification of 6His-tag Fabs was carried out as follows: two weeks post-transfection, antibodies were affinity purified from about 175 mL of ExpiCHO medium (ThermoFisher) cleared by centrifugation and filtration on a 2 mL CaptureSelect LC-kappa (mur) affinity column (Thermo Scientific).

The Fab was eluted with 0.1M Glycine, pH 3.4, neutralized with Tris pH 8.8 and applied to a HisPur Ni-NTA resin (Thermo Scientific) column. The Fab was eluted with 250 mM imidazole and buffer exchanged with PD-10 columns (Cytiva) to 50 mM Tris pH 8.0, 0.15M NaCl and 3mM CaCl_2_. To obtain a single pure species of 19A11 for crystallography, a minor glycosylated product (10% of the total) was removed by incubating with ConA slurry (GE Healthcare) for 4 hours at 4°C. on a rotator. For production of a single species of 66E8 for crystallography, 6 μM kifunensine was added to the ExpiCHO culture media at the time of transfection. For production of a single species of 66E8 for crystallography, 6 μM kifunensine was added to the ExpiCHO culture media at the time of transfection and Endo Hf (NEB) treatment of LC-kappa purified protein (∼10,000 U/mg protein at 4°C for 3 hours) was done prior to HisPur Ni-NTA purification. Isolation of a single species for each Fab was verified by PAGE and activation of cellular E-cadherin was confirmed by Colo205 activation assay (described below). Purification of StrepTag Fabs from ExpiCHO culture medium was performed using StrepTactin XT Superflow High Capacity resin (IBA), elution with 50 mM biotin, followed by buffer exchange with PD-10 columns to 50 mM Tris pH 8.0, 0.15 M NaCl and 3 mM CaCl_2_.

#### E-cadherin EC1-5/19A11 complex formation

hEC1-5 was incubated with a 1.6x molar excess of conA-purified 19A11-6His Fab and incubated overnight at 4°C. Complex was purified with SEC with a Superose 6 10/300 GL column and concentrated to 11.5 mg/mL in 50 mM Tris, 150 mM NaCl, 3 mM CaCl_2_, pH 8.0.

### Nanodisc preparation

Purified full-length E-cadherin was concentrated to 8-10 μM and mixed with the nanodisc scaffolding protein MSP1D1 (Sigma) at a 5 fold molar excess. Samples later bound to 46H7 and 19A11 were incubated with catenins, as detailed in a previous study^38^. 100mM DMPC / 200 mM CHAPS in 20 mM Tris 7.4, 100 mM NaCl was added to a final DMPC/CHAPS concentration of 8 mM /16 mM, respectively. The final ratio for disc formation was 1 E-cadherin : 5 MSP1D1 : 80 DMPC per disc. This mixture was incubated for 30 min at 20°C, then 0.8 g/mL Amberlite® XAD®-2 (Sigma-Aldrich 10357) was added to remove detergent and incubated for a further 2 hours at 20°C. Assembled E-cadherin-TwinStrep discs were purified away from empty discs with a 1 mL StrepTactin XT column equilibrated in BB. Column was washed with BB and eluted with BB + 50 mM Biotin. E-cadherin nanodiscs were further purified with SEC using a Superose 6 10/300 GL SEC column (GE). Peak fractions containing all components were collected, and glycerol was added to 2.5%. Protein was then concentrated to 0.2-0.4 mg/mL, flash frozen, and stored at -80°C.

### X-ray crystallography

#### Crystallization

The hEC1-2/66E8 complex was crystallized at 10.4 mg/mL at 14C and mixed 1:1 with a solution of 12.5% (w/v) PEG 4000, 20% (v/v) 1,2,6-hexanetriol, 0.1M GlyGly/AMPD pH 8.5, and 0.03M of each lithium sulfate, sodium sulfate, and potassium sulfate (Morpheus II A10). The hEC1-5/19A11 complex was crystallized at 11.5 mg/mL at 14C and mixed 2:1 with a solution of 0.1 M sodium HEPES pH 7.0 and 15% w/v PEG 4000 (ProPlex B11). Upon harvesting, crystals were cryocooled in liquid nitrogen. hE-cad1-2/66E8 crystals did not require additional cryoprotection. hE-cad EC1-5/19A11 crystals were dipped in a 15% ethylene glycol solution prior to cryocooling.

#### Data collection and processing

X-ray diffraction data for both complexes were collected at the LS-CAT beamline 21-ID- F at the Advanced Photon Source. Data were collected at 100K at a wavelength of 0.97872 Å. All data were integrated and scaled using XDS and XSCALE.

#### Structure solution and refinement

Structures were solved by molecular replacement using Phaser within the CCP4 program suite. Each structure utilized a model for each the cadherin and antibody: PDB entries 2o72 and 2v17, respectively (hEC1-2/66E8); PDB entries 3q2v and 6cxy, respectively (hEC1-5/19A11). Structures were refined in iterative cycles of real space refinement in Coot and reciprocal space refinement in Phenix. The quality of each model was assessed using MolProbity as implemented in Phenix. Final hEC1-2/66E8 structure was deposited to the PDB as 6VEL. Final hEC1-5/19A11 structure was deposited as PDB 7STZ. Structure refinement data are provided in Table 4.

**Table 1.** Recombinant functional antibody fragments used in this study

| Antibody | Effect on E-cad | Epitope |
| --- | --- | --- |
| 19A11 | Activating | EC1 |
| 59D2 | Activating | EC1 |
| 66E8 | Activating | EC1-2 |
| 46H7 | Neutral | EC3 |
| 67G8 | Blocking | EC5 |

**Table 2.** PDBs created or referenced in this study

| PDB | Protein |
| --- | --- |
| 3Q2V | Mouse E-cadherin EC1-5 |
| 2O72 | Human E-cadherin EC1-2 |
| 6CXY | Human E-cadherin EC1-2/19A11 Fab |
| 7STZ | Human E-cadherin EC1-5/19A11 Fab |
| 6VEL | Human E-cadherin EC1-2/66E8 Fab |

**Table 3.**
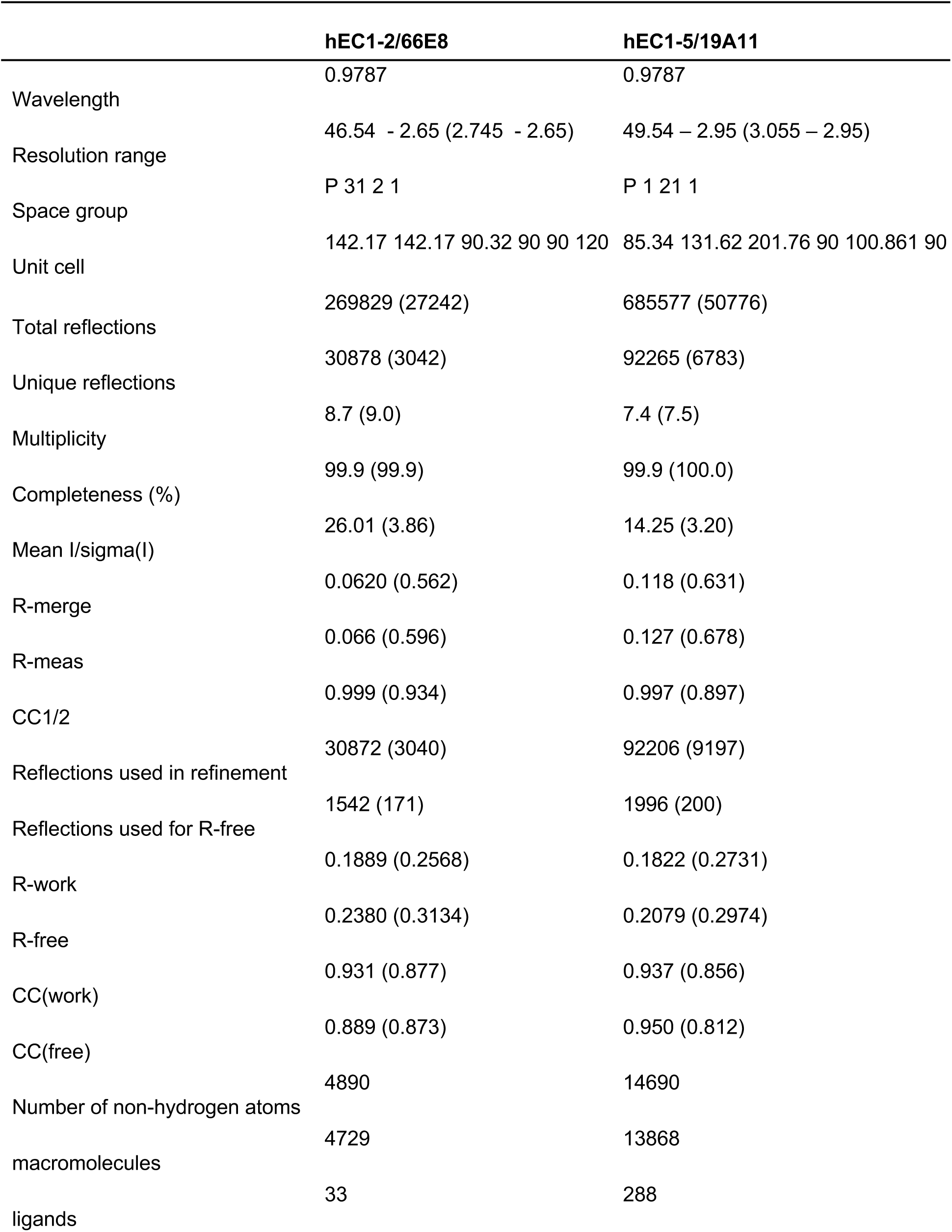
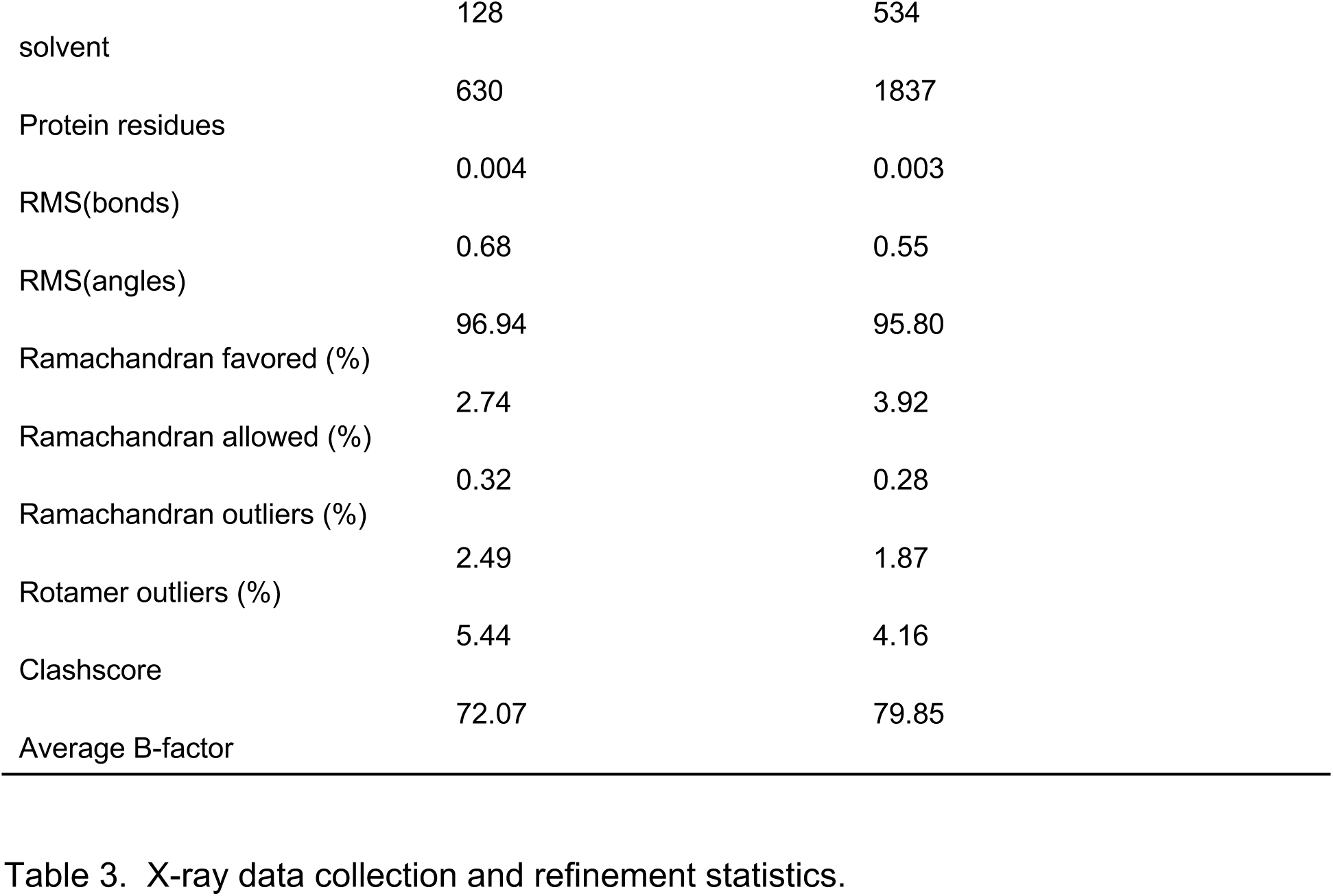
X-ray data collection and refinement statistics.

**Table 4.** Cryo-EM data collection, reconstruction, and refinement.

| Cryo-EM data collection and processing statistics |  |  |  |  |
| --- | --- | --- | --- | --- |
| Sample | Full-length E-cadherin-catenin complex + 19A11 | Full-length E-cadherin-catenin complex + 46H7 | Full-length E-cadherin + 59D2 | Full-length E-cadherin + 67G8 |
| **Data collection** |  |  |  |  |
| Microscope | Titan Krios | Titan Krios | Titan Krios | Titan Krios |
| Voltage (kV) | 300 | 300 | 300 | 300 |
| Magnification | 130000x | 130000x | 105000x | 105000x |
| Detector | Gatan K2 | Gatan K2 | Gatan K3 | Gatan K3 |
| Data collection software | Leginon | Leginon | Leginon | Leginon |
| Electron exposure ($e^-/\text{\AA}^2$) | 40 | 40 | 47 | 64 |
| Defocus Range ($\mu\text{m}$) | -1 - -2.5 | -1 - -2.5 | -1 - -2.5 | -1 - -2.5 |
| Pixel size ($\text{\AA}$) | 0.525 | 0.525 | 0.84 | 0.42 |
| **Data processing** |  |  |  |  |
| Number of micrographs | 1823 | 2004 | 3805 | 1655 |
| Final particle images | 99879 | 67509 | 331400 | 97712 |
| Symmetry imposed | C1 | C1 | C1 | C1 |
| Map resolution ($\text{\AA}$) 0.143 FSC threshold | 4.85 | 4.75 | 6.24 | 5.55 |

### Cryo-EM sample preparation and data collection

For full-length hE-Cadherin-catenin nanodiscs (19A11 + 46H7), 10 μL nanodiscs at 0.2 mg/mL (2 μg) were incubated with 2 μL of 1 mg/mL (2 μg) Fab for 1 hour at 20°C. After incubation, these were diluted in half, and 3 μL diluted sample was applied to a glow discharged C-Flat™ Holey Carbon Grid CF-2/2-4C, 400 mesh Cu (Electron Microscopy Sciences CF-224C-50). This was incubated for 1 min, then blotted using a Vitrobot Mark IV (FEI) at 4°C, 100% humidity, 4-5 sec blot time, 0 blot force, then plunge frozen in liquid ethane. Data was collected on a 300 kV Titan Krios G3 with a K2 Summit camera in super-resolution mode (0.525 Å/pix).

For full-length hE-Cadherin-only nanodiscs with Fab (59D2 and 67G8), 10 μL nanodiscs at 0.2 mg/mL (2 μg) were incubated with 2 μL of 1 mg/mL (2 μg) Fab for 1 hour at 20°C. After incubation, these were diluted to 1/3, and 3 μL diluted sample was applied to a glow discharged Au-Flat 2/2 200 Gold Mesh grid (AUFT222-50) (59D2) or C-Flat™ Holey Carbon Grid CF-2/2-4C, 400 mesh Cu (67G8). This was incubated for 1 min, then blotted using a Vitrobot Mark IV (FEI) at 4°C, 100% humidity, 4-5 sec blot time, 0 blot force, then plunge frozen in liquid ethane. Data was collected on a 300 kV Titan Krios G3 with a K3 Summit camera. 67G8 data was collected in super-resolution mode (0.42 Å/pix); 59D2 data was collected in regular counting mode (0.84 Å /pix).

For full-length hE-Cadherin-only nanodiscs examined only as 2D averages (WT, W2A, K14E, WT+19A11, W2A+19A11), 10 μL nanodiscs at 0.2 mg/mL (2 μg) were incubated with 2 μL of 1 mg/mL (2 μg) Fab, if applicable, for 1 hour at 20°C. After incubation, or in samples with no Fab, immediately, these were diluted in half, and 3 μL diluted sample was applied to a glow discharged C-Flat™ Holey Carbon Grid CF-2/2- 4C, 400 mesh Cu (Electron Microscopy Sciences CF-224C-50). This was incubated for 1 min, then blotted using a Vitrobot Mark IV (FEI) at 4°C, 100% humidity, 4-5 sec blot time, 0 blot force, then plunge frozen in liquid ethane. Data was collected on a 200 kV Glacios Cryo-TEM with a K2 Summit camera at 1.16 Å /pix.

All datasets were queued and collected using Leginon ^47^.

We note again that the data here for 19A11 and 46H7 grids were collected with full cadherin-catenin complex. As the results were the same whether or not catenins were bound, as detailed in previous work^38^, subsequent analyses (67G8, 59D2, no fab, mutants) were done on just nanodisc-embedded FL-hE-cadherin-TwinStrep with no intracellular proteins bound.

### Cryo-EM data processing

For FL-hE-cadherin-catenin complex ND + 19A11Fab, data for 1823 movies were aligned with MotionCor2 in Relion 3.0.3^48^ then CTF was estimated with CryoSPARC v2.14^49^. 205,013 particles were picked with a crYOLO v1.3.6^50^ model trained on this dataset, extracted in Relion 3.0.3, then re-imported back to cryoSPARC for further processing. 204,452 particles were extracted and immediately subjected to ab initio reconstruction into 3 volumes. Classes 0 and 1 (145,965 particles) were selected, and underwent Homogeneous refinement based on the class 0 model in CryoSPARC. These particles then went through a round of 2D averaging to clean out junk particles. The final 99,879 selected particles then were homogeneous refined using the previous refinement reconstruction as a model, then all particles and the model were used for cryoSPARC non-uniform refinement, leading to a gold standard FSC final resolution of 4.85 Å after mask auto-tightening. Although all views were represented, preferred views were more represented in the reconstruction, resulting in a skewed range of directional views. As done in Billesbølle et al.^51^, particle stacks were then exported with csparc2star.py as part of the pyem package^52^; stacks were created in Relion and imported into cisTEM^53^. New half maps were generated with the generate3D module; maps were sharpened in cisTEM. Local resolution was estimated in cryoSPARC, and FSCs were calculated with Relion, showing identical resolution estimates to cryoSPARC’s NU-refinement calculated resolution. Although average resolution was unchanged as measured by Relion, new maps generated in cisTEM showed a less broad range of local resolution estimates, as well as improved 3D FSC (0.966 in the cisTEM generated map vs 0.844 for cryoSPARC) as measured by the 3D FSC server^54^, compared to the raw cryoSPARC NU-refinement reconstructions.

For FL-hE-cadherin-catenin complex ND + 46H7Fab, 2004 movies were imported, motion corrected, and CTF estimated with cryoSPARC 2.14. Poor and low resolution exposures were removed, resulting in 1962 micrographs. A small number of particles were manually picked, 2D averaged, and used as templates for particle picking in cryoSPARC. 531,229 particles were picked and extracted. After 2 rounds of 2D averaging to remove bad and broken particles, as well as unbound Fabs, the remaining 108,022 particles were inputted to Ab initio reconstruction in cryoSPARC with 4 models. These 4 models then went through Heterogeneous refinement, also in cryoSPARC. Two classes (0+1) were picked, resulting in 67,509 final particles that underwent homogeneous refinement (using class 1 as the model), then non-uniform refinement, resulting in a final reconstruction at 4.75 Å resolution by gold-standard FSC. As described previously, new maps were created in cisTEM with the generate3D command, local resolution was estimated in cryoSPARC, and overall resolution FSCs were generated with Relion.

For FL-hE-cadherin ND + 59D2 Fab, 3805 movies were imported, patch motion corrected, and patch CTF estimated with cryoSPARC 2.14. Template picker was used to pick 1,077,980 particles; after exposure curation, 830,126 particles were extracted and underwent 3 rounds of 2D averaging to remove junk particles, leaving 534,216 particles. 100,000 of these underwent Ab initio reconstruction into 3 classes; all 534,216 particles were then heterogeneously refined to these 3 classes. Classes 0 and 1 (331,400 particles) underwent homogeneous refinement, then non-uniform refinement with class 0 as the starting model, still in cryoSPARC. These were re-extracted at 640 bin 2 box sizes, then went through one additional round of non-uniform refinement, resulting in a 6.24 A reconstruction. As described previously, new maps were created in cisTEM with the generate3D command, local resolution was estimated in cryoSPARC, and overall resolution FSCs were generated with Relion.

For FL-hE-cadherin ND + 67G8 Fab, 1213 movies at 0 degree tilt and 442 movies at 30 degree tilt from 2 data collections were separately patch motion corrected and CTF estimated with cryoSPARC 2.14. Each then had particles picked in crYOLO (0: 279009; 30: 77977) and went through two rounds of 2D averaging leading to a final particle count of 192133 particles. The combined particles went through one final round of 2D averaging, leading to a particle count of 116371 particles. These then went into an Ab initio reconstruction of 4 classes, followed by heterogeneous refinement of these 4 class models. Classes 0+1+3 (97712 particles) were selected and homogeneous refined, followed by non-uniform refinement, leading to a final 3D reconstruction at 5.55 A resolution by gold-standard FSC. As described previously, new maps were created in cisTEM with the generate3D command, local resolution was estimated in cryoSPARC, and overall resolution FSCs were generated with Relion.

FL-hE-cadherin ND WT, W2A, K14E, WT+19A11, W2A+19A11 all went through analogous data analysis procedures to ensure comparative results. Each of these datasets was also repeated a second time with fresh sample to verify repeatability.

Briefly, movies were aligned with Patch Motion correction with CryoSPARC v2.14; CTF was estimated with CryoSPARC Patch CTF. Particles were then picked using crYOLO, using a model trained on WT E-cadherin, extracted in Relion, and re-imported back into cryoSPARC, where they were extracted with 512 bin 4 box sizes and subjected to two rounds of 2D classification to weed out junk particles. A third round of classification where the initial classification uncertainty factor was set to 6, and 40 iterations of classification were performed, was used to separate different dimer conformations.

### Bio-Layer interferometry

BLI kinetics assays were performed on an Octet Red96 at 23°C, shaking at 1000 rpm. Protein was diluted in kinetics buffer: 50 mM Tris pH 8.0, 150 mM NaCl, 3mM CaCl_2_, 0.25 mg/mL BSA, 0.005% Tween-20. Ni-NTA sensors (ForteBio) were equilibrated for 60 seconds, then E-cadherin EC1-5-TwinStrep-8His was loaded onto the sensor for 180 seconds, followed by another 60 second baseline. Sensors were then immersed into a 1:3 dilution series of anti-E-cadherin Fabs in kinetics buffer until they reached desired concentrations, then dipped into empty kinetics buffer to determine off-rates. ForteBio data analysis software was used to calculate kinetics parameters such as *k_on_*, *k_off_*, and K_D_ using a 1:1 binding global fit model. Assays were repeated at least twice with different Fab preparations to ensure consistent results. For 19A11 and 46H7, both ficin-cleaved untagged Fabs and TwinStrep tagged Fabs were tested; both showed similar affinities regardless of presence of tag (Extended Data Figure 4F).

### Analytical size exclusion chromatography

hE-cadherin EC1-5 TwinStrep constructs were incubated with 3.2x molar excess Fab (2x by mass) at 4°C for ∼16 hours. Mixtures were then injected into a Superose 6 10/300 column. For analysis, elution times were multiplied by the 0.5 mL/min flow rate to calculate elution volume in mL. Fractions were run on 5-20% SDS-PAGE gels to examine protein composition of each peak.

### Colo205 Activation Assay

The Colo205 activation assay was performed as described previously^17^. Briefly, Colo205 cells were densely plated on 96-well plates precoated with 0.1μg/mL rat-tail collagen (Sigma-Aldrich) overnight and treated with activating concentrations of Fabs for 5 hours. Activation was determined by the extent of a morphological change from round cells with distinct borders to a compact epithelial appearance and loss of obvious cell borders.

## Data Availability Statement

X-ray crystallographic structures generated during the current study are available in the Protein Data Bank with accession codes 7STZ (hE-cadherin EC1-5/19A11) and 6VEL (hE-cadherin EC1-2/66E8). Cryo-EM density maps are available in the Electron Microscopy Data Bank with accession codes 25883 (hE-cadherin/19A11), 25884 (hE-cadherin/46H7), 25886 (hE-cadherin/59D2), and 25892 (hE-cadherin/67G8). All other datasets generated and/or analyzed during the current study are available from the corresponding author on reasonable request.

## Acknowledgements

We thank Justin Kollman and Melody Campbell for valuable discussions and input. We thank Joel Quispe, Quinton Beedle, and Sasha Dickinson for training and support on cryo-EM equipment. We thank also David Veesler and Kelly Lee for use and training on the Octet Red96. Cryo-EM data was analyzed with the Cybertron high-performance computing cluster at Seattle Children’s Research Institute. Cryo-EM was performed at the University of Washington Arnold and Mabel Beckman cryo-EM center.

---

**Table S1.** Individual measurements and kinetics calculations of Fab affinity.

|  | **19A11** | **46H7** | **59D2** | **67G8** | **66E8** |
| --- | --- | --- | --- | --- | --- |
| K<sub>D</sub> calc | 8.05E-09 | 9.46E-07 | 1.19E-12 | 9.59E-09 | 9.01E-08 |
|  | 5.10E-09 | 2.58E-07 | 2.42E-12 | 4.56E-09 | 9.65E-08 |
|  | 6.71E-09 | 3.24E-07 | 1.81E-12 | 1.09E-08 | 9.18E-08 |
|  | 6.42E-09 |  | 1.88E-12 | 9.52E-09 |  |
| Average | 6.57E-09 | 5.09E-07 | 1.83E-12 | 8.64E-09 | 9.28E-08 |
| Std dev | 1.21015E-09 | 3.79602E-07 | 5.0349E-13 | 2.79469E-09 | 3.33593E-09 |

**Extended Data Figure 1.**
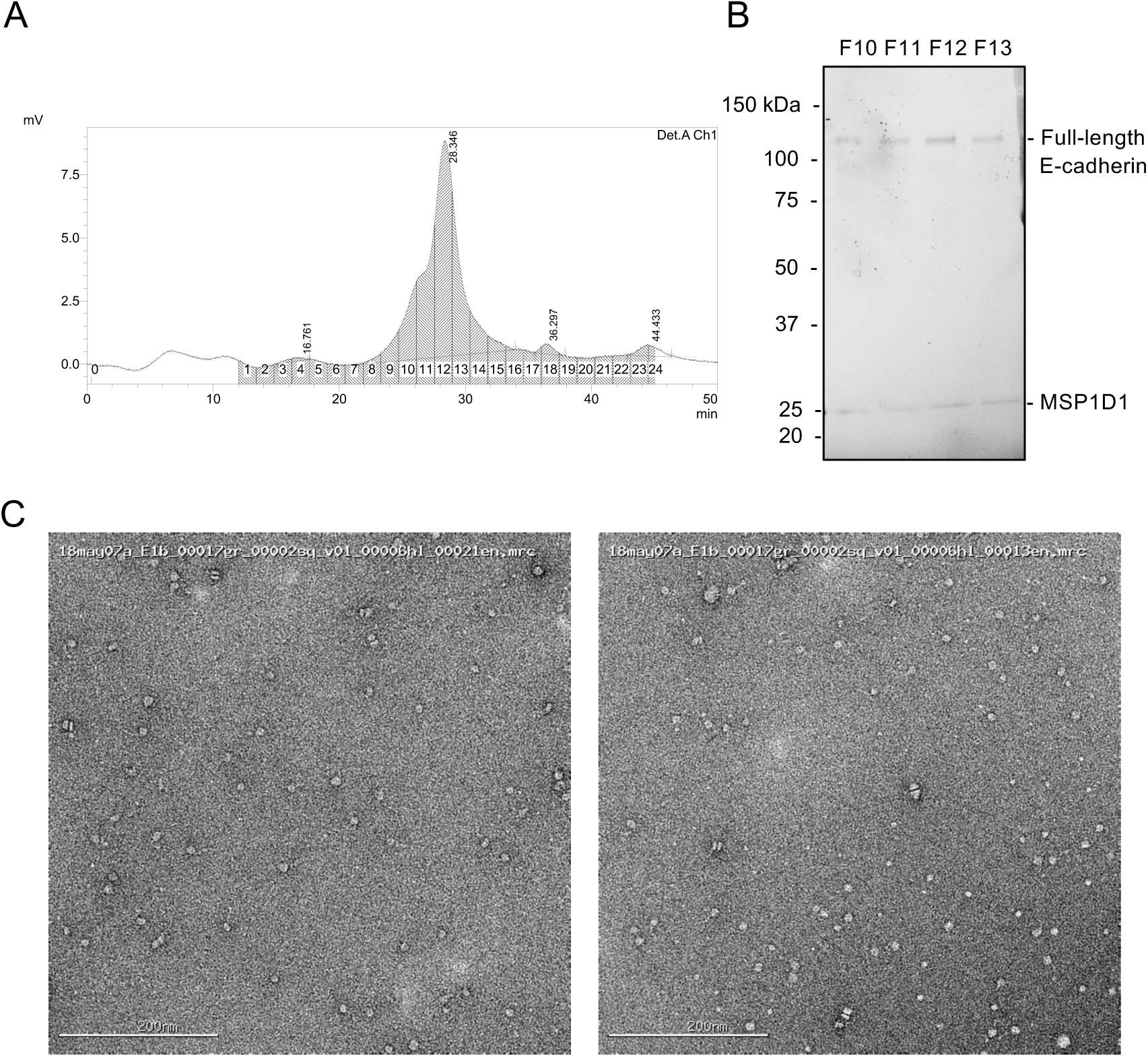
Full-length E-cadherin is embedded in MSP1D1 nanodiscs. (A) SEC chromatogram of FL-hE-cadherin nanodiscs. (B) SDS-PAGE gel of SEC peak fractions indicating presence of E-cadherin and MSP1D1 membrane scaffold protein (C) Negative stain EM micrographs of E-cadherin nanodiscs. Minor stacking is evident from the calcium content in the buffer.

**Extended Data Figure 2.**
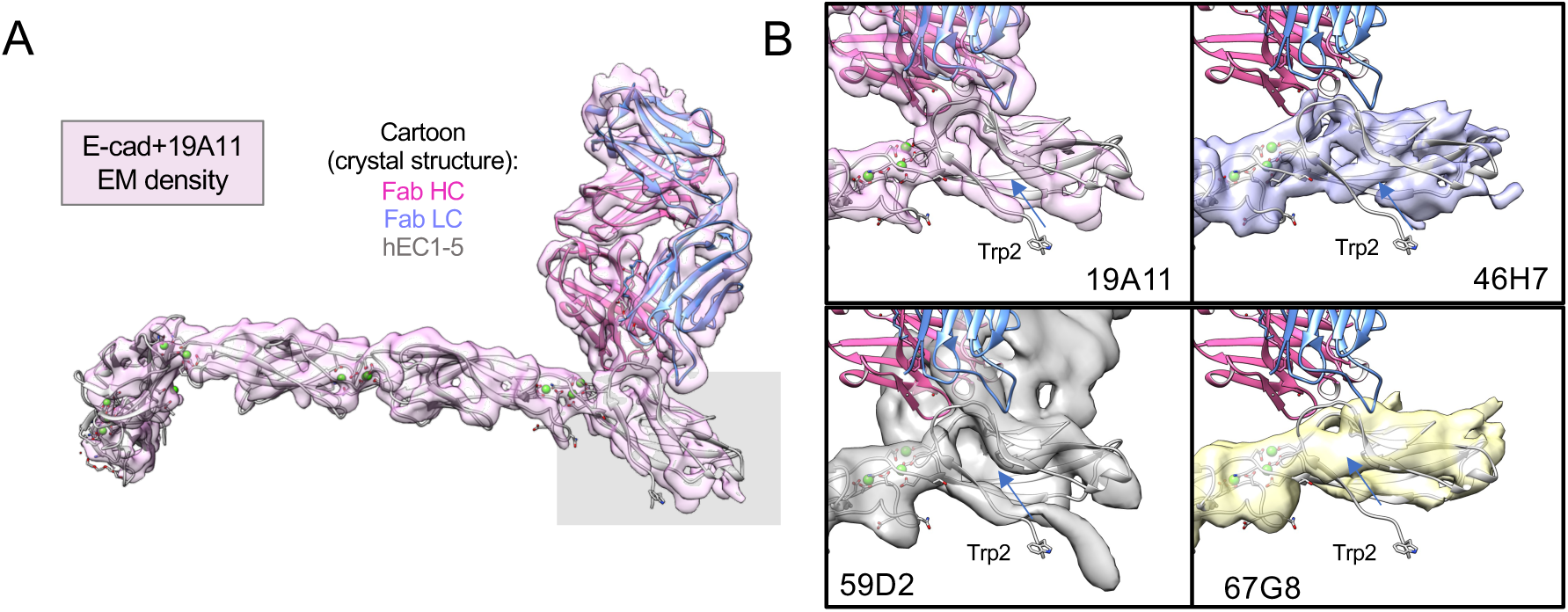
EM reconstructions of activating and non-activating Fabs have variations in EC1 density. (A) Overlay of monomeric EM reconstruction and crystal structure of hEC1-5/19A11. Grey box highlights general EC1 region examined in (B). Closeups of EC1 with each Fab bound, overlaid with EC1-5/19A11 structure to indicate location of beta strand and Trp2. The arrow indicates the location of the hydrophobic pocket.

**Extended Data Figure 3.**
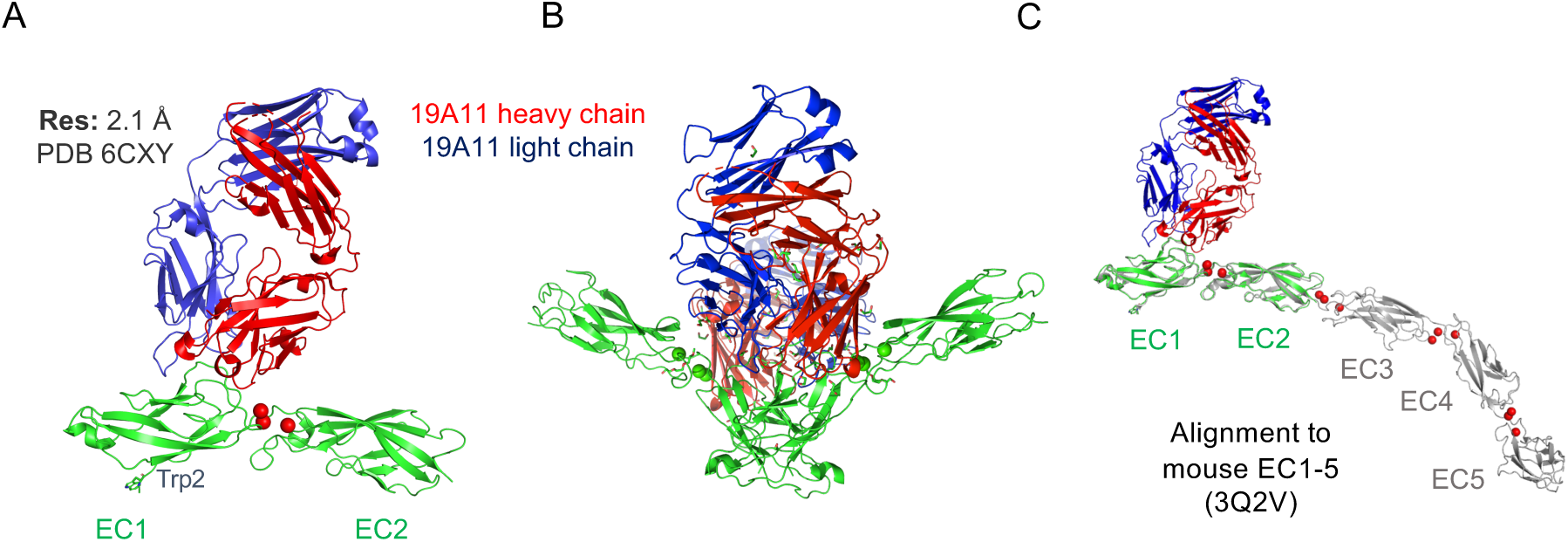
Crystal structure of hEC1-2/19A11 activating Fab. (A) Asymmetric unit of crystal structure indicating Fab epitope in EC1. (B) Strand-swap dimer seen in crystal expansion (C) Overlay with mouse EC1-5 PDB indicating epitope location in full ectodomain.

**Extended Data Figure 4.**
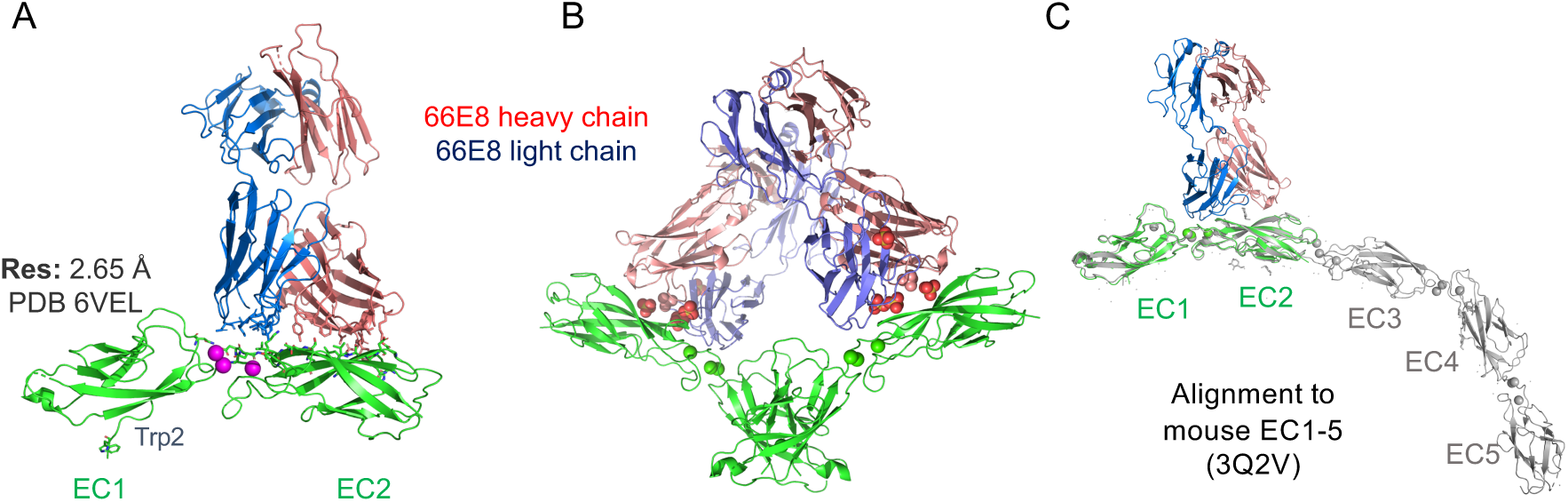
Crystal structure of hEC1-2/66E8 activating Fab. (A) Asymmetric unit of crystal structure indicating Fab epitope in EC2 and the EC1-2 Ca binding site. (B) Strand-swap dimer seen in crystal expansion (C) Overlay with mouse EC1-5 PDB indicating epitope location in full ectodomain.

**Extended Data Figure 5.**
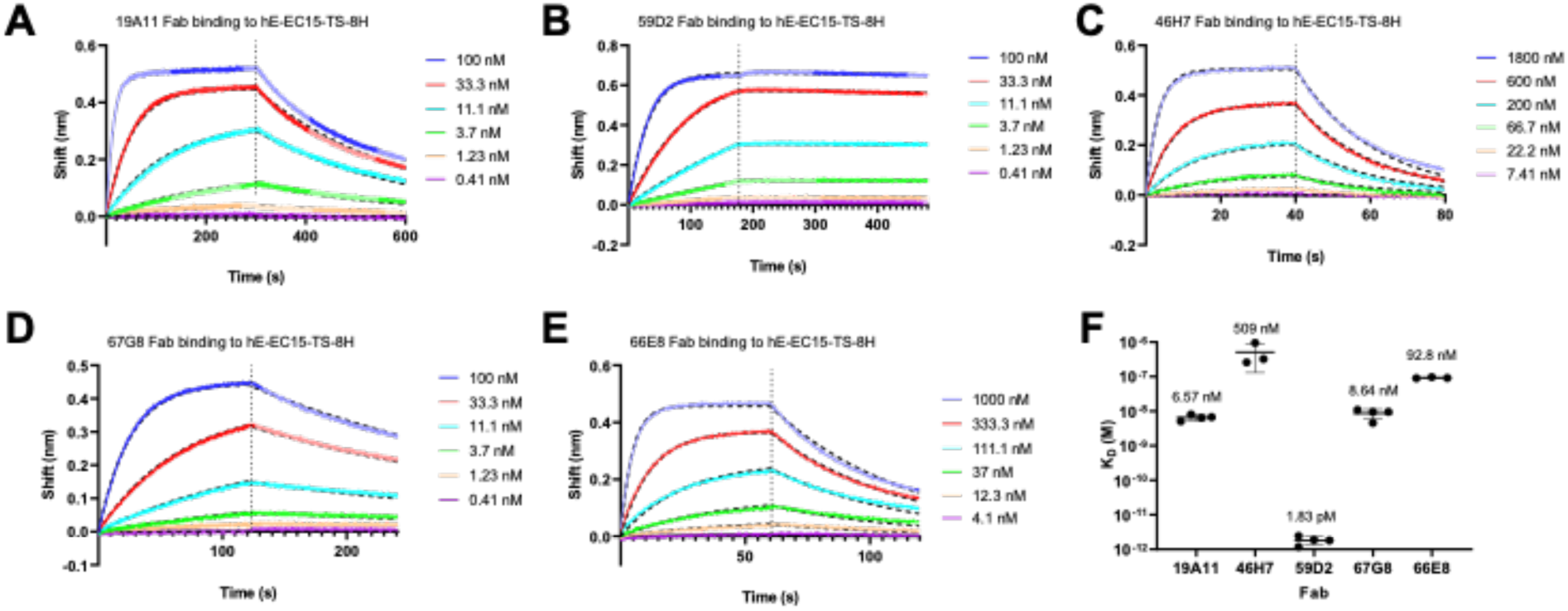
BLI kinetics of Fabs binding to hEC1-5-TS-8His. (A) 19A11 Fab binding curve. (B) 59D2 Fab binding curve. (C) 46H7 binding curve. (D) 67G8 binding curve.(E) 66E8 binding curve (F) summary of individual measurements. Mean K_D_ labeled for each Fab.

**Extended Data Figure 6.**
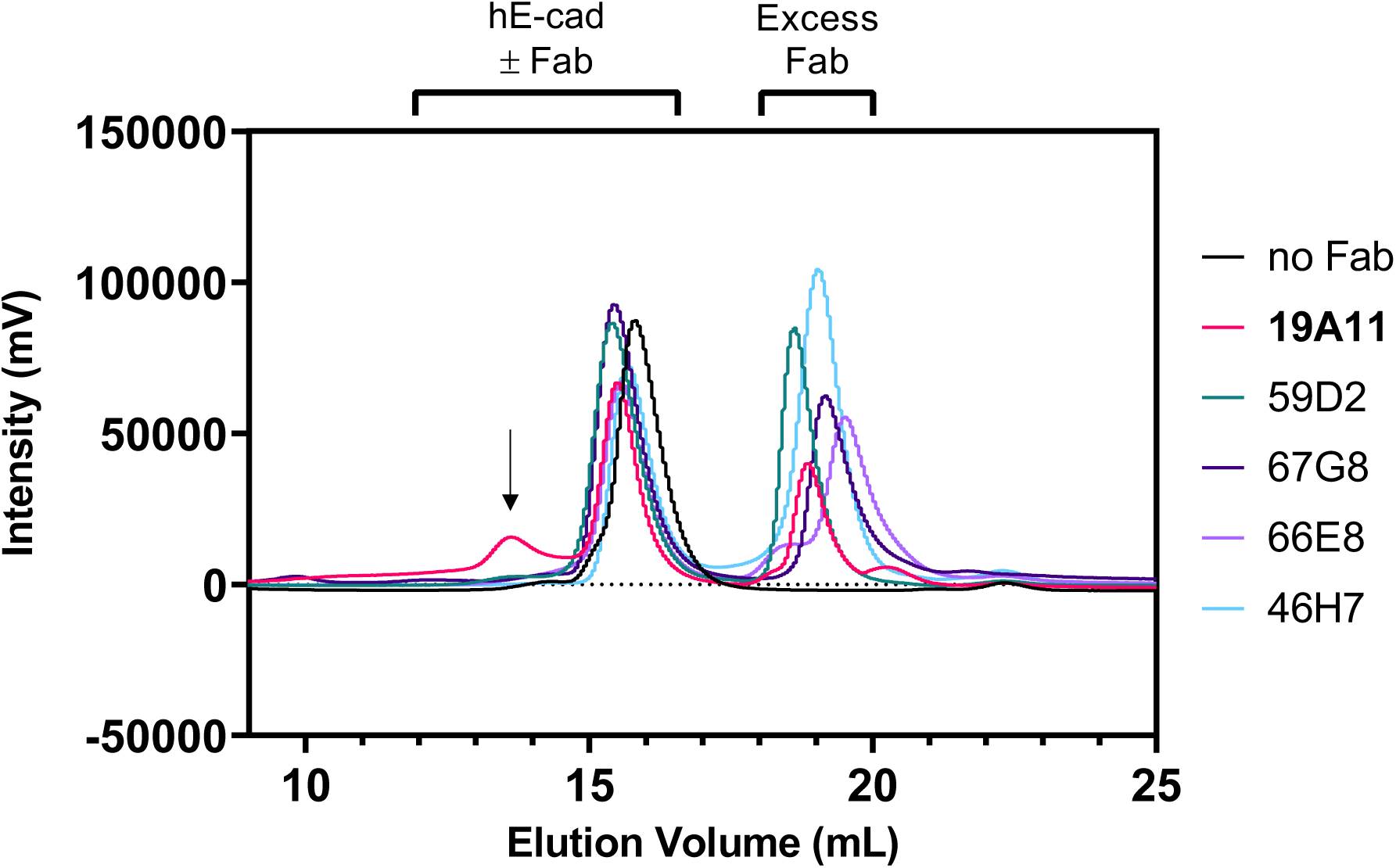
SEC of all recombinant functional antibodies bound to hEC1-5 shows that 19A11 is prominent in its formation of a hEC1-5 dimer peak.

**Extended Data Figure 7.**
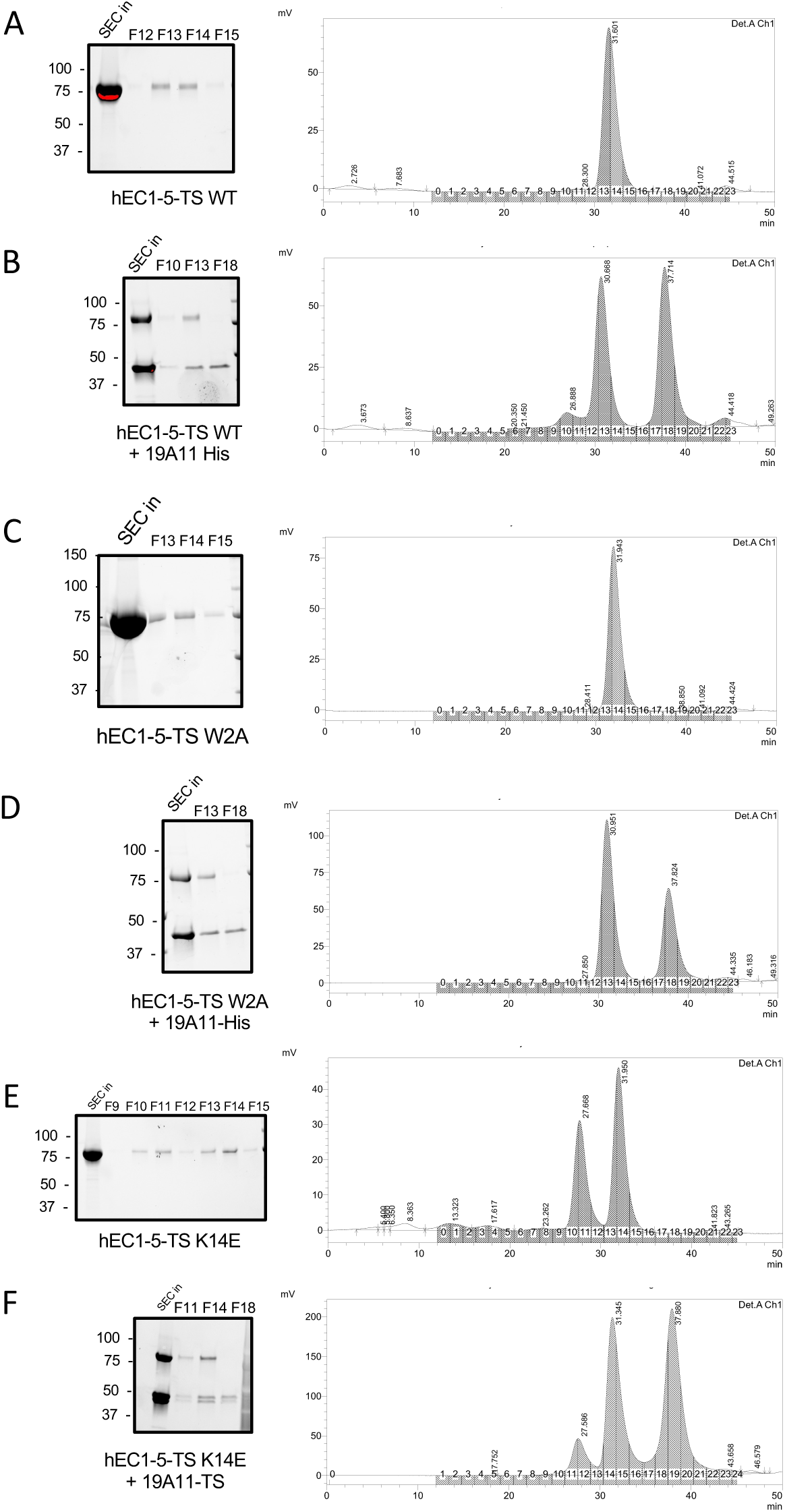
Individual raw SEC chromatograms and gels of fractions of human E-cadherin ectodomain Twin Strep (hEC1-5-TS) mutants bound to 19A11 Fab. (A) hEC1-5-TS WT alone (B) hEC1-5 TS WT mixed with and excess of 19A11 Fab. (C) hEC1-5 TS W2A strand-swap deficient mutant. (D) hEC1-5 TS W2A mixed with and excess of 19A11 Fab. (E) hEC1-5 TS K14E X-dimer blocking mutant alone. (F) hEC1-5 TS K14E mixed with an excess of 19A11 Fab.

**Extended Data Figure 8.**
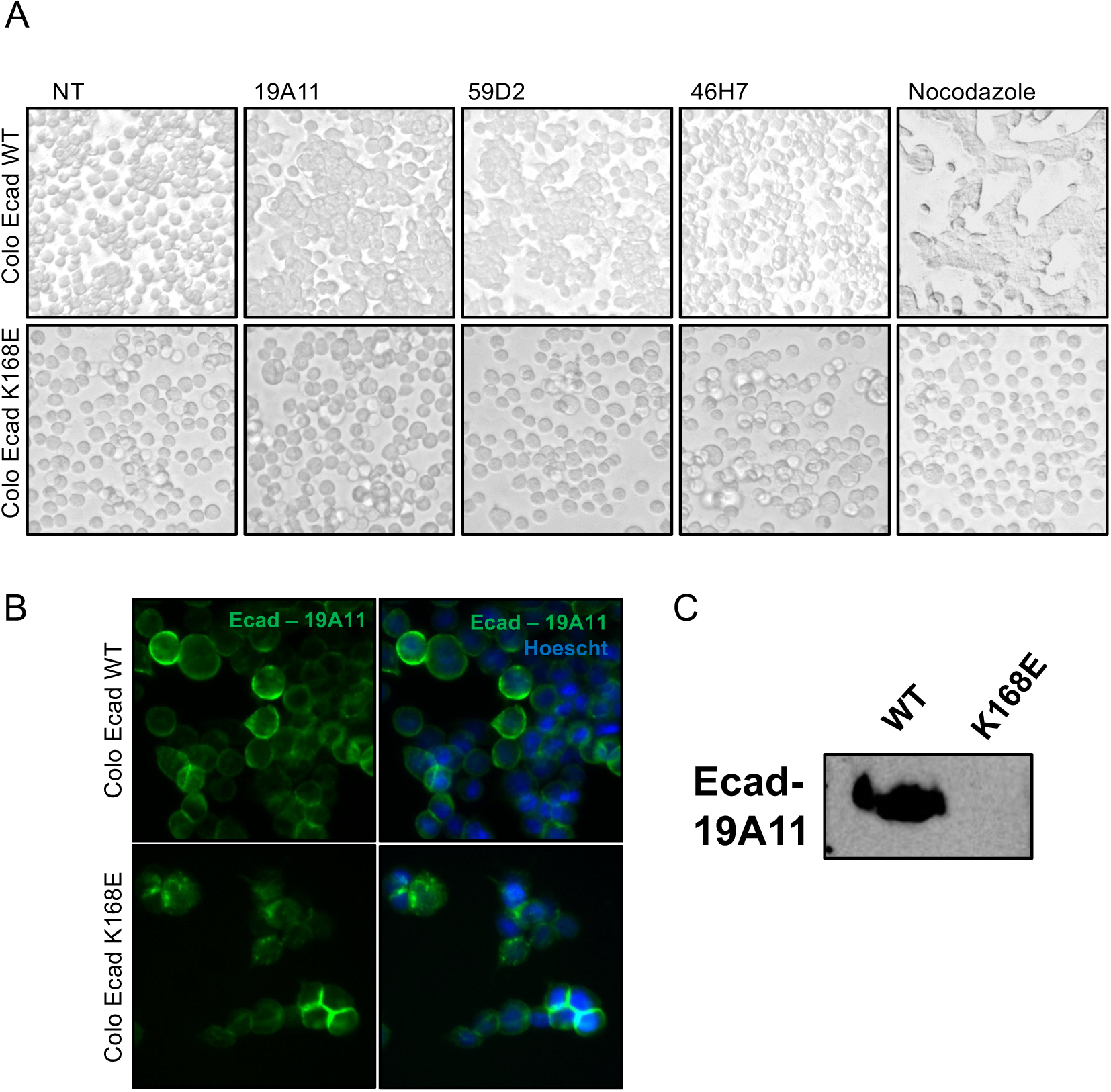
Colo205 activation with K14E/K168E E-cadherin is not rescued by 19A11. (A) Colo205 activation assay of WT E-cadherin expressing cells and K168E E-cadherin cells with full mAb treatment. NT = no treatment. (B) Immunofluorescence staining of WT and K168E E-cadherin Colo205 cells with 19A11 full mAb. (C) Western blot of Colo205 cell lysates expressing either WT E-cadherin or K168E.

