## Supplementary Figures for "Regulation of multiple dimeric states of E-cadherin by adhesion activating antibodies revealed through Cryo-EM and X-ray crystallography"

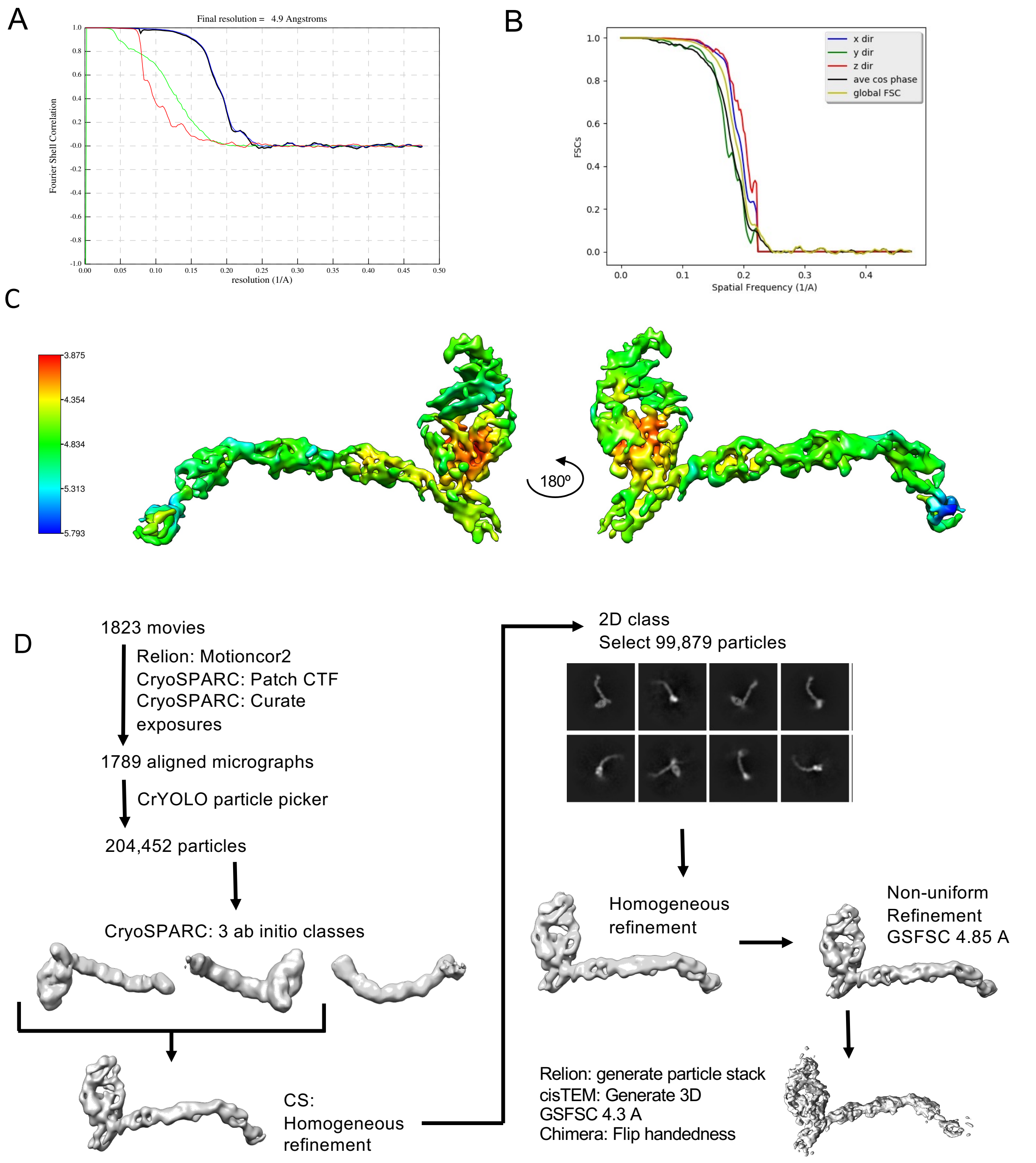


Figure S1. Cryo-EM characterization of FL-hE-cadherin + 19A11Fab. **(A)** Relion Gold-Standard 0.143 FSC resolution of final reconstruction. **(B)** Directional resolution of map calculated by 3DFSC server. Calculated sphericity: 0.966. **(C)** Local resolution estimation over sharpened final 3D reconstruction. **(D)** Data processing pipeline toward final reconstruction.

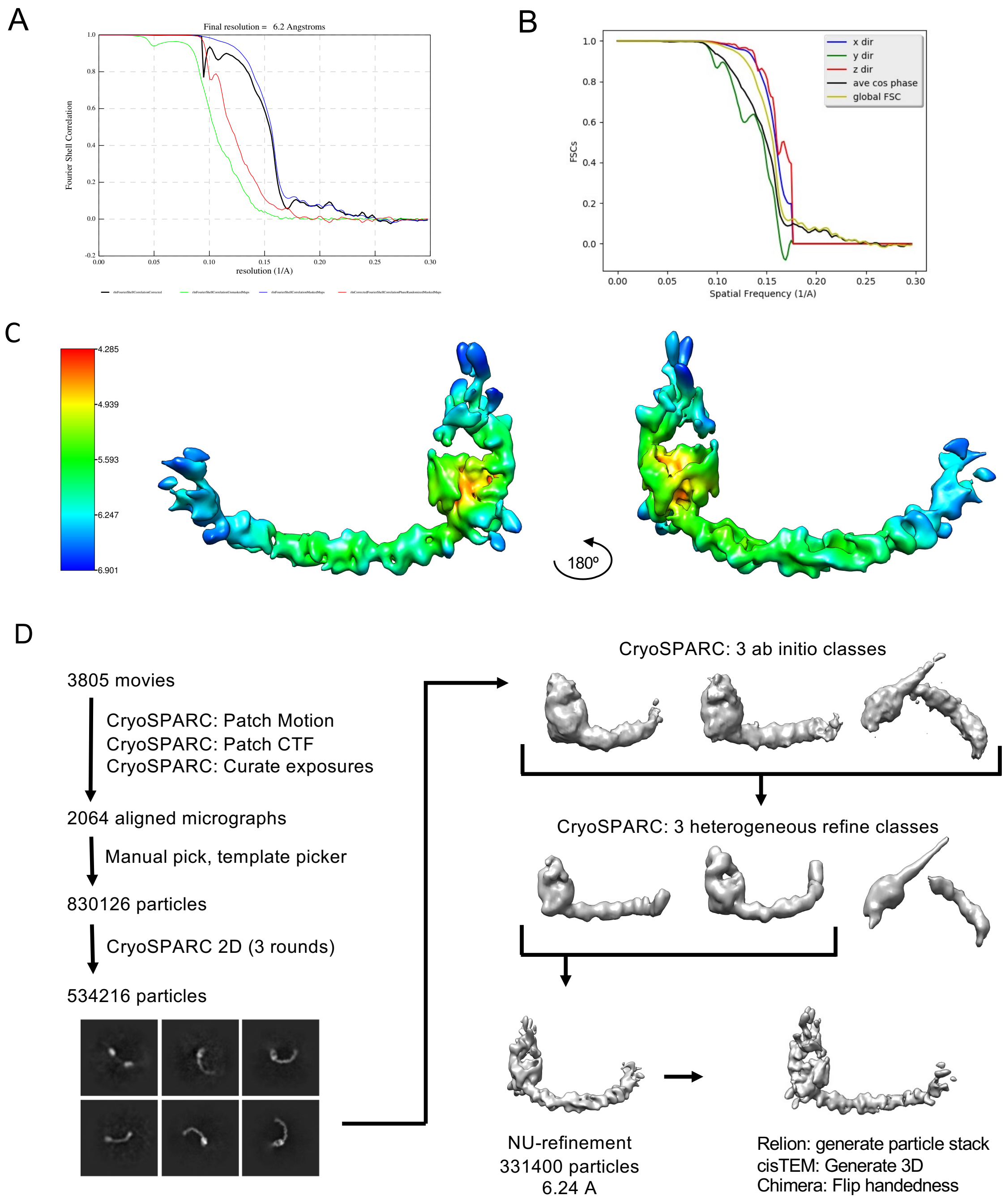


Figure S2. Cryo-EM characterization of FL-hE-cadherin + 59D2Fab. (**A)** Relion Gold-Standard 0.143 FSC resolution of final reconstruction. **(B)** Directional resolution of map calculated by 3DFSC server. Calculated sphericity: 0.973. **(C)** Local resolution estimation over sharpened final 3D reconstruction. **(D)** Data processing pipeline toward final reconstruction.

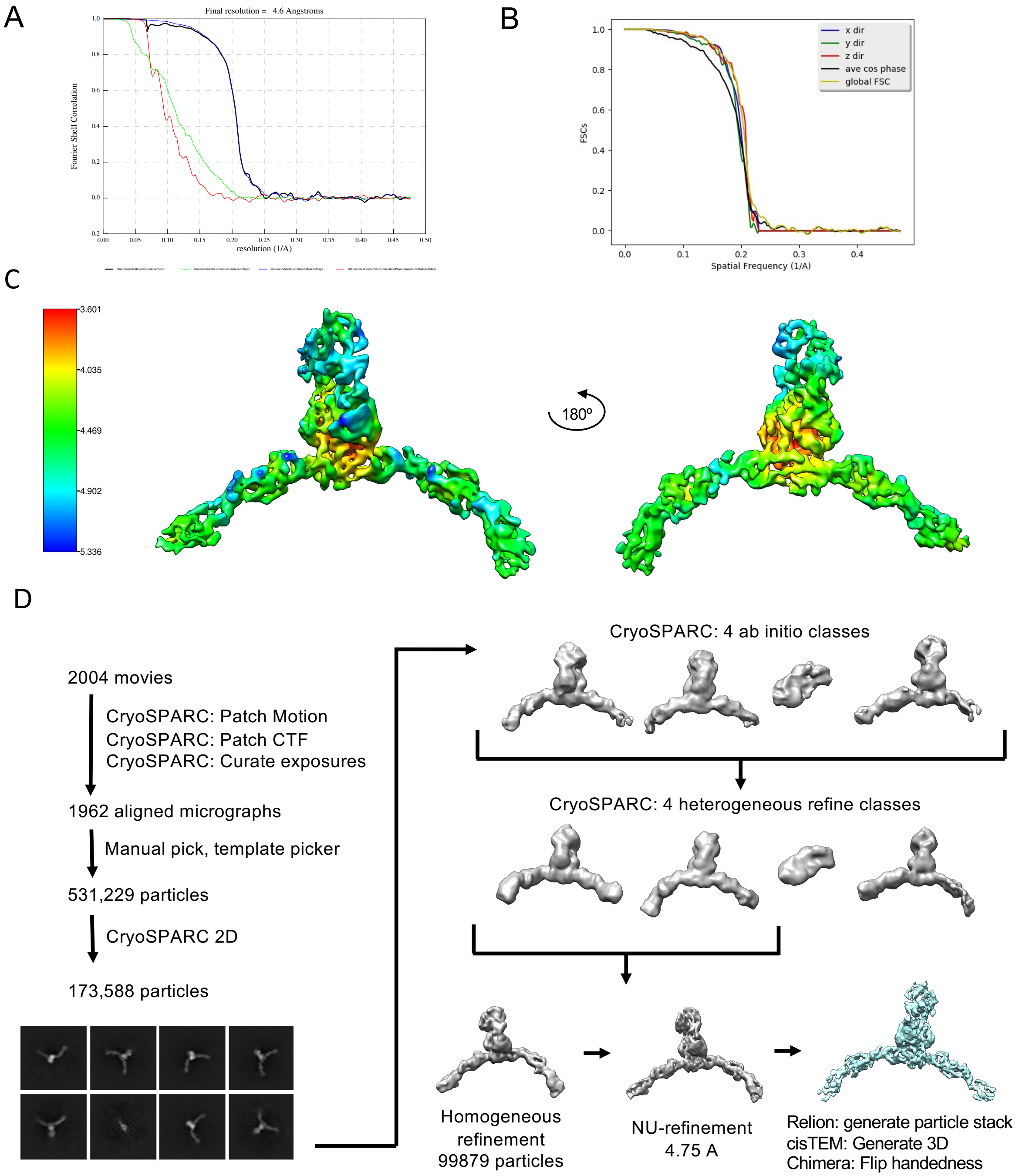


Figure S3. Cryo-EM characterization of FL-hE-cadherin + 46H7Fab. **(A)** Relion Gold-Standard 0.143 FSC resolution of final reconstruction. **(B)** Directional resolution of map calculated by 3DFSC server. Calculated sphericity: 0.982. **(C)** Local resolution estimation over sharpened final 3D reconstruction. **(D)** Data processing pipeline toward final reconstruction.

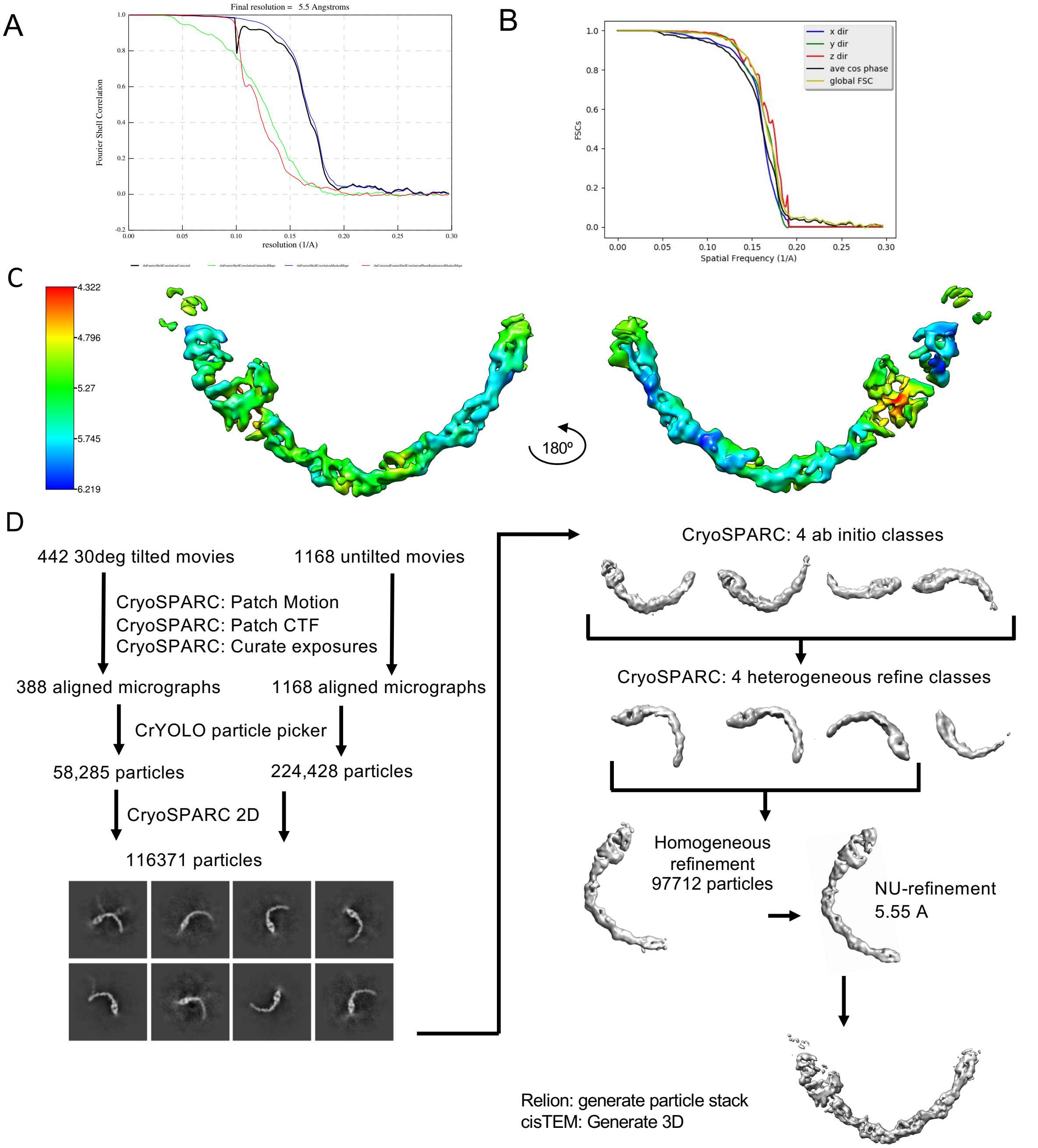


Figure S4. Cryo-EM characterization of FL-hE-cadherin + 67G8Fab. **(A)** Relion Gold-Standard 0.143 FSC resolution of final reconstruction. **(B)** Directional resolution of map calculated by 3DFSC server. Calculated sphericity: 0.963. **(C)** Local resolution estimation over sharpened final 3D reconstruction. **(D)** Data processing pipeline toward final reconstruction.
